# Integrative genomics identifies lncRNA regulatory networks across 1,044 pediatric leukemias and extra-cranial solid tumors

**DOI:** 10.1101/2020.12.10.420257

**Authors:** Apexa Modi, Gonzalo Lopez, Karina L. Conkrite, Chun Su, Tsz Ching Leung, Sathvik Ramanan, Elisabetta Manduchi, Matthew E. Johnson, Daphne Cheung, Samantha Gadd, Jinghui Zhang, Malcolm A. Smith, Jaime M. Guidry Auvil, Daniela S. Gerhard, Soheil Meshinchi, Elizabeth J. Perlman, Stephen P. Hunger, John M. Maris, Andrew D. Wells, Struan F.A. Grant, Sharon J. Diskin

## Abstract

Long non-coding RNAs (lncRNAs) play an important role in gene regulation and contribute to tumorigenesis. While pan-cancer studies of lncRNA expression have been performed for adult malignancies, the lncRNA landscape across pediatric cancers remains largely uncharted. Here, we curate RNA sequencing data for 1,044 pediatric leukemia and solid tumors and integrate paired tumor whole genome sequencing and epigenetic data in relevant cell line models to explore lncRNA expression, regulation, and association with cancer. We report a total of 2,657 robustly expressed lncRNAs across six pediatric cancers, including 1,142 exhibiting histotype-specific expression. DNA copy number alterations contributed to lncRNA dysregulation at a proportion comparable to protein coding genes. Application of a multi-dimensional framework to identify and prioritize lncRNAs impacting gene networks revealed that lncRNAs dysregulated in pediatric cancer are associated with proliferation, metabolism, and DNA damage hallmarks. Analysis of upstream regulation via cell-type specific transcription factors further implicated distinct histotype-specific and developmental lncRNAs. We integrated our analyses to prioritize lncRNAs for experimental validation and showed that silencing of *TBX2-AS1*, our top-prioritized neuroblastoma-specific lncRNA, resulted in significant growth inhibition of neuroblastoma cells, confirming our computational predictions. Taken together, these data provide a comprehensive characterization of lncRNA regulation and function in pediatric cancers and pave the way for future mechanistic studies.

---

Long non-coding RNAs (lncRNAs) are transcribed RNA molecules greater than 200 nucleotides in length that do not code for proteins. These molecules account for 70% of the expressed human transcriptome and provide a key aspect of gene regulation^1–4^. Compared to protein coding genes (PCGs), lncRNAs typically have fewer exons, weaker conservation, and lower abundance^3^. Despite this, lncRNAs have been shown to play significant roles in both transcriptional and post-transcriptional gene regulation^5^. LncRNAs perform these roles by physically interacting with a variety of substrates, including proteins (transcription co-factors), RNAs (microRNA sponges), and DNA (chromatin interaction scaffolds)^1,2,6,7^. While the mechanisms and function for the majority of lncRNAs remain unknown^3,8^, those that have been experimentally characterized are involved in a variety of cellular processes^6^ including gene silencing (*ANRIL*)^9^, modulation of chromatin architecture (*Xist*)^10^, and pre-mRNA processing (*MALAT1*)^11^. LncRNAs are also important in development^12^. For example, the *H19* lncRNA is involved in imprinting^13^, while the well-conserved *TUNA* lncRNA controls stem cell pluripotency and lineage differentiation^14^.

Dysregulation of lncRNA expression has been widely observed in cancer^3,15,16^ and studies have shown that lncRNAs play important roles in tumor initiation and progression^17^. LncRNAs can function as tumor suppressors, such as the *PANDA* lncRNA which regulates DNA damage response in diffuse large B-cell lymphoma^18^; however, many more lncRNAs appear to be oncogenes. Examples include the *HOTAIR* and *PVT1* lncRNAs which promote proliferation in various cancers through tissue specific mechanisms^19,20^. Pan-cancer analyses of lncRNA expression in adult malignancies have uncovered many cancer-associated lncRNAs^3,15–17,21,22^. Identification of functional lncRNAs amongst the large set of cancer-associated lncRNAs, however, remains challenging^15,23^. Current methods to identify putative functional lncRNAs involve identifying lncRNA-specific genetic aberrations^15,16,24^ or using lncRNA expression to predict overall patient survival^16^. To more systematically address how lncRNAs drive the pathogenesis of cancer, recent computational methods seek to assign function to these molecules based on predicted target genes and regulatory network models. These methods have been applied to adult malignancies and allow for more focused hypotheses to be tested^21,22^.

LncRNA studies and evidence of related function in pediatric cancers have been primarily limited to neuroblastoma (NBL)^25–30^, T-lymphoblastic leukemia (T-ALL)^31,32^, and more recently glioblastoma^33^. *CASC15* and *NBAT-1* are a sense-antisense lncRNA pair that map to a NBL susceptibility locus identified by genome-wide association study^26,34^. Both lncRNAs are downregulated in high-risk NBL tumors and have been shown to be involved in cell proliferation and differentiation^25,26^. In pediatric T-ALL, the NOTCH-regulated lncRNA, *LUNAR1*, promotes T-ALL cell growth by sustaining IGF1 signaling^32^. To date, it is unknown whether lncRNAs function as common drivers across multiple pediatric cancers, or if instead, the majority of lncRNAs influence oncogenesis in a histotype-specific manner. Furthermore, given that pediatric cancers typically arise from primitive embryonic and mesodermal cells, rather than adult epithelial cells, it is unclear whether adult cancer lncRNA drivers will also be implicated in childhood cancer.

Here, we perform a pan-pediatric cancer study of lncRNAs across 1,044 pediatric leukemias and extra-cranial solid tumors^35,36^. We present the landscape of lncRNA expression across these childhood cancers and perform integrative multi-omic analyses to assess tissue specificity, regulation, and putative function. To validate our approach, we show that silencing of the top-prioritized NBL-specific lncRNA, *TBX2-AS1*, impairs NBL cell growth in human-derived NBL cell line models.

## Results

### The lncRNA landscape of pediatric cancers

To define the repertoire of lncRNAs expressed in childhood cancers, we analyzed RNA-sequencing data from six distinct pediatric cancer histotypes profiled through the Therapeutically Applicable Research to Generate Effective Treatments (TARGET) project (https://ocg.cancer.gov/programs/target/data-matrix) (**Online Methods; Supplementary Table 1).** This curated set of 1,044 leukemia and solid tumor samples includes 280 acute myeloid leukemia (AML), 190 B-lymphoblastic leukemias (B-ALL), 244 T-lymphoblastic leukemias (T-ALL), 121 Wilms tumors (WT), 48 extracranial rhabdoid tumors (RT), and 161 neuroblastomas (NBL) (**Fig. 1a**). Since one of our goals was to identify novel cancer-associated lncRNAs, we performed guided *de novo* transcriptome assembly using StringTie v1.3.3^37^ with the GENCODE v19 database ^38^ as a gene annotation reference (**Supplementary Fig. 1**). Expressed gene sequences that did not match exons and transcript structures of any known gene in the GENCODE v19 or RefSeq v74 databases were considered putative novel genes (**Supplementary Fig. 1, Online Methods**). Of these novel genes, we identified candidate lncRNAs by using the PLEK v1 algorithm^39^ to assess non-coding potential, and then additionally filtered hits by transcript length, exon read coverage, and genomic location (**Fig. 1a, Online Methods, Supplementary Fig. 1**). As validation of our lncRNA discovery pipeline, we observed that 36% (87 of 242) of identified novel lncRNAs not annotated in Gencode v19 (hg19) were indeed annotated in the more recent Gencode v29 (hg38) genome build (**Supplementary Table 2**). To ensure that we selected robustly expressed genes in the setting of cancer heterogeneity and sequencing variability, we applied a conservative expression cutoff of Fragments Per Kilobase of transcript per Million mapped reads (FPKM) >1 in at least 20% of samples for each cancer. Across all cancers there were 15,588 PCGs, 2,512 known lncRNAs, and 145 novel lncRNAs expressed, though the total number of expressed genes varied per cancer (**Fig 1b, Supplementary Table 3**). Principal component analysis (PCA) of lncRNA gene expression showed that blood (AML, B-ALL, T-ALL) and solid (NBL, WT, RT) cancers form two distinct groups. Moreover, individual cancer histotypes clustered more closely using lncRNA expression than PCG expression alone (**Supplementary Fig. 2a-b**), consistent with the known tissue specific nature of lncRNA expression and function^3^.

**Fig 1:**
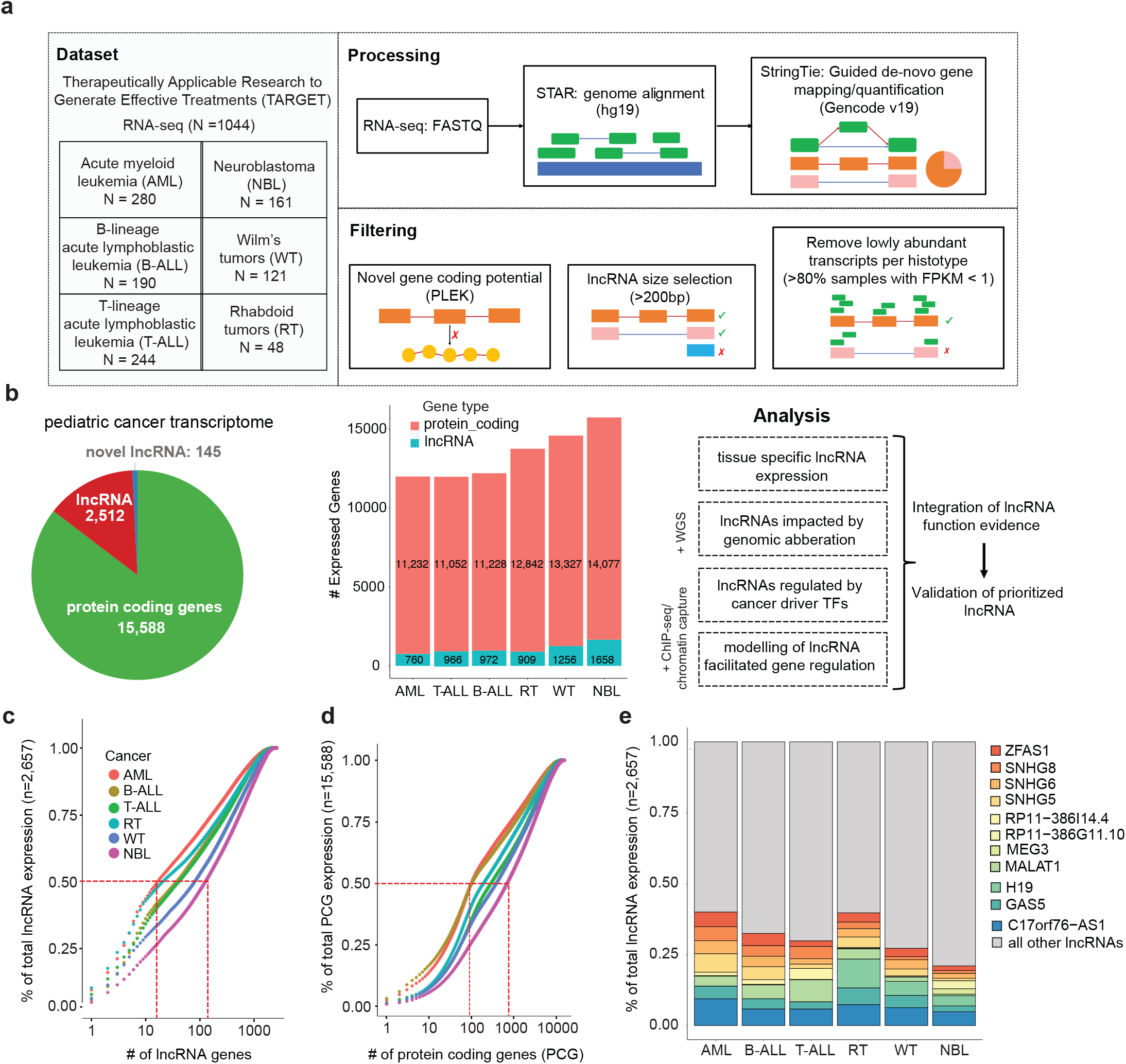
Pan-pediatric transcriptome characterization. **a.** Overview of pan-pediatric cancer RNA-seq dataset and schematic of data processing and filtering. Reads from RNA-seq fastq files were aligned using the STAR algorithm and then gene transcripts were mapped in a guided *de novo* manner and quantified via the StringTie algorithm. Genes were considered novel if they did not have transcript exon structures matching genes in the GENCODE v19 or RefSeq v74 databases. Novel genes were assigned as lncRNAs based on length >200bp and non-coding potential calculated using the PLEK algorithm. Transcripts with low expression (FPKM <1 in >80% samples) were not considered for further analysis. **b.** Pie graph showing the quantity of expressed and robustly expressed protein coding genes, GENCODE/RefSeq annotated lncRNAs, and novel lncRNAs. High confidence expressed genes are distinguished from all expressed genes. Adjoining schematic gives overview of additional data types that were integrated with transcriptome data: WGS, ChIP-seq, and chromatin capture. Listed are the analyses used to elucidate lncRNAs with functional roles in pediatric cancer. **c.** Cumulative expression plots comparing the number of lncRNAs and (**d**) protein coding genes, respectively, that constitute the total sum of gene expression (FPKM) per pediatric cancer. **e.** Percentage of total lncRNA expression (FPKM) accounted for by the union of top five expressed lncRNAs per cancer (total 11 lncRNAs).

Overall, lncRNAs had lower average expression compared to PCGs resulting in fewer highly expressed lncRNAs (**Supplementary Fig. 2c**). Between 10-100 (3.7%) lncRNAs accounted for 50% of the total sum of lncRNA expression (**Fig. 1c**). In contrast, between 100-1000 (6.4%) PCGs accounted for 50% of the total sum of PCG expression (**Fig. 1d**). We examined the union of the top five most highly expressed lncRNAs across pediatric cancers (total 11 lncRNAs). Some of these lncRNAs had higher expression in the blood cancers (*MALAT1* and *RP11-386I14.4*), in the solid cancers (*H19)*, or in only one cancer, such as *MEG3* and *RP11-386G11.10* in NBL (**Fig. 1e**). Five of these lncRNAs were among the top 10 lncRNAs expressed across normal tissues in the Genotype-Tissue Expression (GTEx) project ^40^. Specifically, *C17orf76-AS1 (LRRC75A-AS1), MALAT1, GAS5, SNHG6, SNHG8* were expressed ubiquitously in 30 of the 49 GTEx tissues (**Supplementary Table 4**).

### Tissue specific lncRNA expression distinguishes pediatric cancers

To evaluate more formally the tissue specific expression of lncRNAs, we annotated all genes with a tissue specificity index (tau score)^41,42^ (**Online Methods**). The established tau score ranges from 0 (ubiquitous expression) to 1 (tissue-specific). As an example, the highly expressed lncRNA *C17orf76-AS1* yielded a tau score of 0.296 in this study, indicating ubiquitous expression (**Supplementary Fig. 2d**). In contrast, the highly expressed *MEG3* lncRNA, which is known to have tissue-specific expression in NBL^30,43^, yielded a tau score of 0.986 (**Supplementary Fig. 2e**). Overall, we observed that lncRNAs yielded a higher tau score range and mean, and thus greater tissue specific expression than PCGs (t-test p=1.62×10^−42^). Novel lncRNAs had the greatest tissue specific expression (t-test: vs proteins- p=1.62×10^−42^, vs known lncRNAs- p = 3.39×10^−13^) (**Fig. 2a**). A tau score threshold of 0.8 has been suggested to distinguish tissue specific genes^42^, and using this cutoff we identified 1,142 (42%) tissue specific (TS) lncRNAs (**Fig. 2b, Supplementary Table 5**). To assess how well TS lncRNAs distinguish cancers, we performed clustering based on the top five highest expressed TS lncRNAs per cancer (30 total). The expression of just these lncRNAs was sufficient to cluster samples of the same cancer type (**Fig. 2c**). Furthermore, the blood and solid cancers separately clustered together with little expression overlap observed between the two groups across the 30 genes (**Fig. 2c**). Finally, we identified a similar proportion of TS lncRNAs (38%, n = 1624) across 12 adult cancers from The Cancer Genome Atlas (TCGA) (**Online Methods**) and observed that adult cancer tissue types were also well distinguished based on the expression of the top 5 most TS lncRNAs (**Supplementary Fig. 2f-g**).

**Fig 2:**
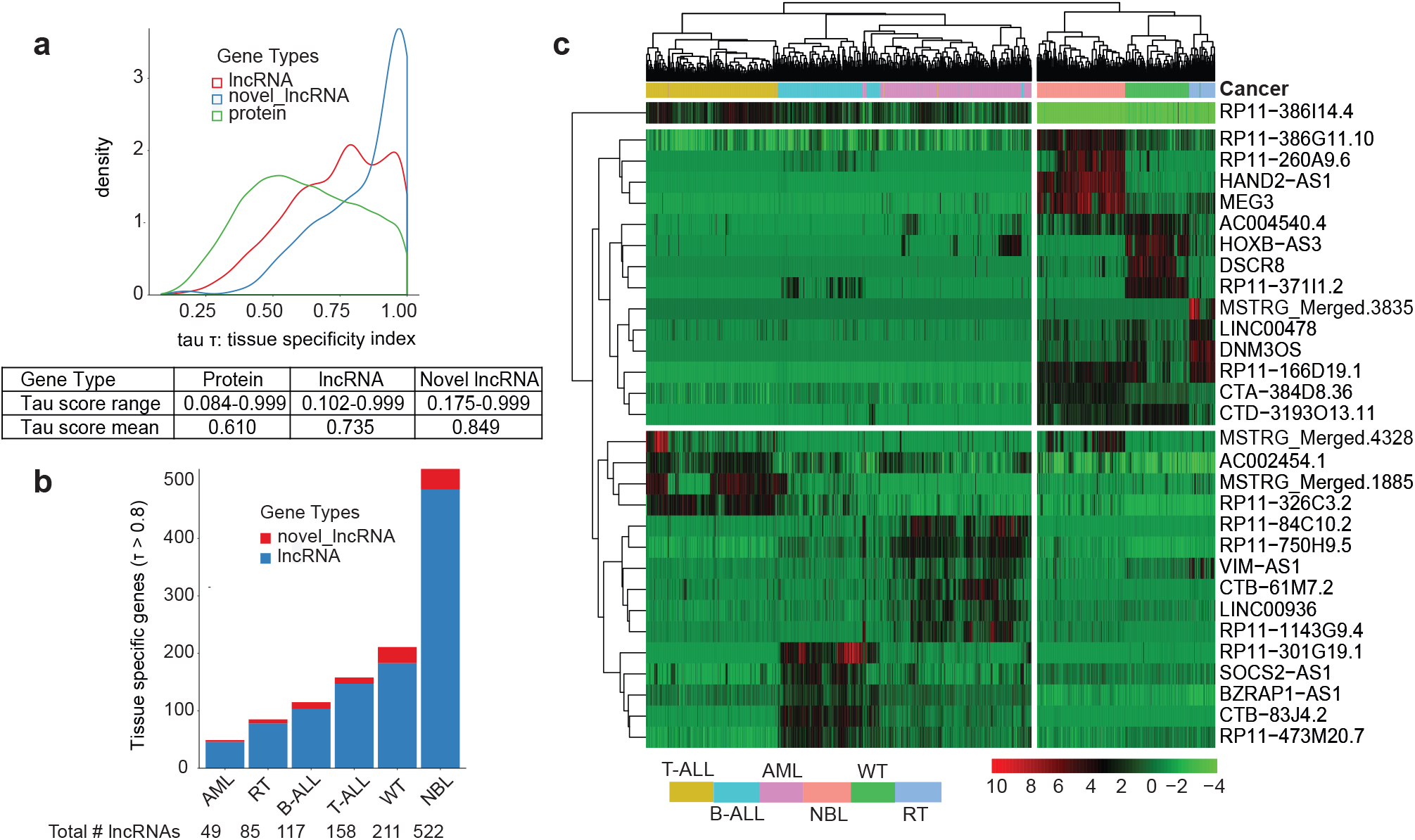
lncRNAs exhibit tissue specific expression that can distinguish cancers. **a.** Tissue specificity index (tau score) which ranges from 0 (ubiquitously expressed) to 1 (tissue specific) is plotted for genes across three gene types: protein coding genes, lncRNAs, and novel lncRNAs. Table shows the tau score range and mean per gene type. **b.** Number of tissue specific known and novel lncRNAs in each cancer as defined by tissue specific gene threshold: tau score > 0.8. **c.** Heatmap showing the hierarchically clustered gene expression for the top five most tissue specific lncRNAs per cancer, ranked by highest tau score. Samples from each cancer cluster together based on expression of these genes alone.

Notably, NBL tumors expressed 2.5x more TS lncRNAs (n=522) than the cancer with the next highest: WT (TS lncRNAs: n=211), and 10x more than AML, which had the least number of TS lncRNAs (n=49) (**Fig. 2b**). To validate NBL’s striking quantity of TS lncRNAs, we first assessed whether immune and stromal cell infiltration^36^ could be contributing to the variety of lncRNAs expressed. We ran the ESTIMATE algorithm as previously described^36^ (**Online Methods**) to determine levels of immune and stromal cell presence in each tumor sample using expression data. Using these purity estimates, we re-calculated each cancer’s tau score and restricted our analysis to NBL samples with either 80% or 90% purity. In both cases, we found that NBL still had the greatest number of TS lncRNAs (n =588 – NBL 90% purity) compared to other cancers (**Supplementary Table 6**). Finally, given that the TARGET NBL RNA-seq dataset is un-stranded, we validated our findings using stranded RNA-seq data in an independent NBL cohort generated through the Gabriela Miller Kids First (GMKF) program (n=223). We observed that 48% of expressed lncRNAs were tissue specific in the GMKF cohort, an increase from the 31% observed in the TARGET cohort (**Supplementary Table 6**). These results confirm lncRNA abundance in NBL and demonstrate that the tau score robustly identifies TS lncRNAs across varying datasets.

### Somatic DNA copy number alterations impact lncRNA expression

Many pediatric cancers are marked by a lower single nucleotide variant (SNV) and insertion-deletion (indel) burden than observed in adult cancers^36^. Instead, large chromosomal events, such as somatic copy number aberrations (SCNAs) and other structural variants (SVs) have been shown to dysregulate protein coding driver genes^36,44^. However, the extent to which large chromosomal alterations impact lncRNAs in pediatric cancers remains unknown. We thus sought to identify SCNAs and SVs using whole genome sequencing (WGS) data from the TARGET project available for NBL (n=146), B-ALL (n=302), AML (n=297), and WT (n=81) (**Online Methods**). We observed that NBL had the greatest frequency of copy number events (**Supplementary Fig. 3a**). The GISTIC v2 algorithm^45^ was applied to detect regions of recurrent SCNA (q-value < 0.25). We identified 673 expressed lncRNAs overlapping 176 significant SCNA regions across the cancers (**Supplementary Table 7**). WGS samples with matched RNA-sequencing were then used to compare lncRNA expression in samples with or without an SCNA event and determine significant differential expression (DE) (**Online Methods**, **Supplementary Table 8**). Across all cancers, between 10-30% of expressed genes overlapping SCNA regions showed significant differential expression based on SCNA, a proportion that was similar for both PCGs and lncRNAs (**Fig 3a**). Altogether, there were 198 (29%) unique lncRNAs with significant DE due to SCNA (**Supplementary Fig 3b**). The majority of the significantly dysregulated lncRNAs were identified in the two cancers with the greatest overall number of expressed lncRNAs, NBL and WT, and mapped to regions with highly recurrent SCNAs in those cancers (chromosomes 1, 7, 11, and 17) (**Fig 3b**).

**Fig 3:**
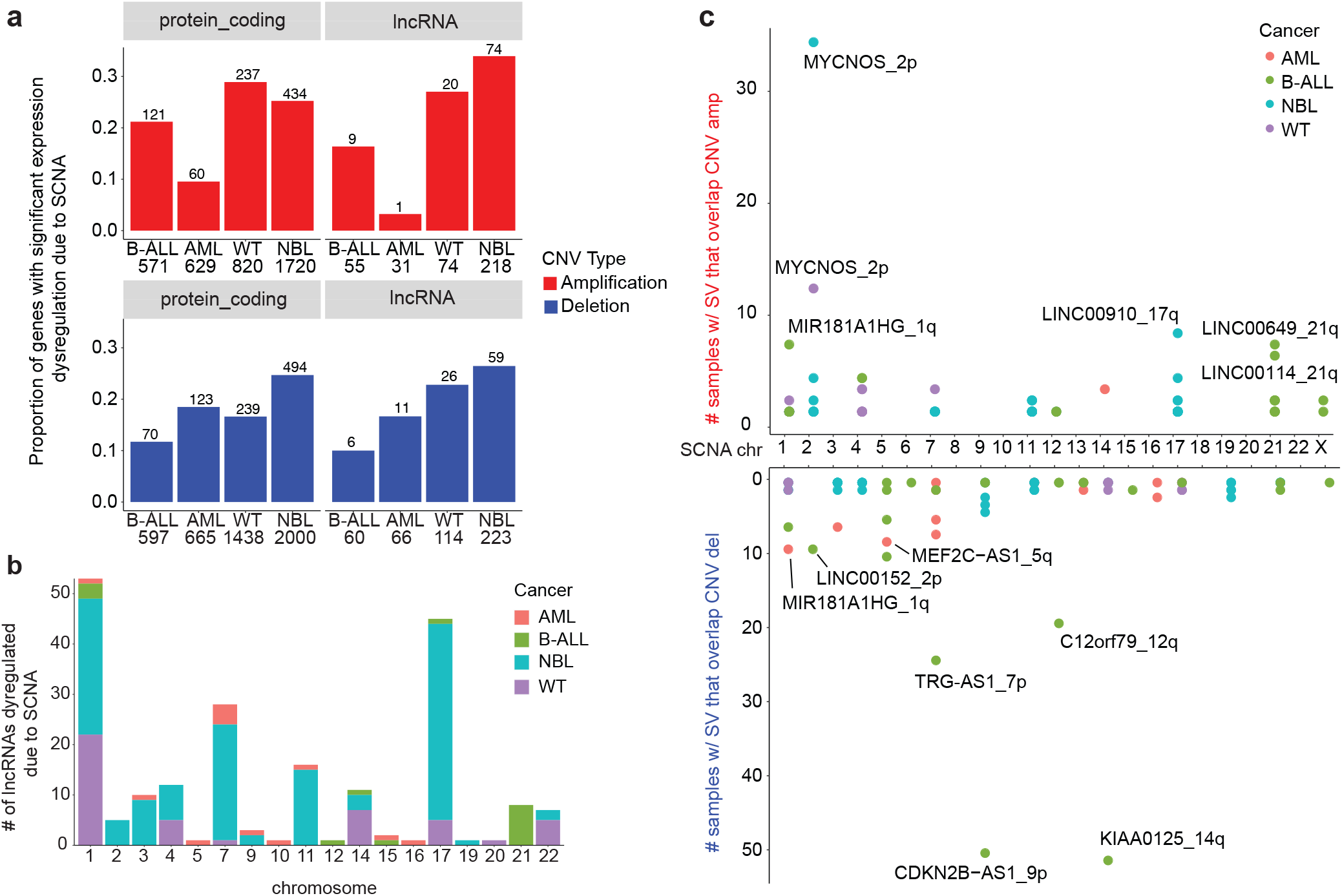
A similar proportion of lncRNAs and protein coding genes are dysregulated due to SCNA. **a.** The proportion of protein coding and lncRNA genes that have significant differential expression due SCNA, separated by copy number type (amplification or deletion). The number of genes found in SCNA loci is shown per cancer. Genes were evaluated to have differential expression due to copy number using the Wilcoxon rank sum test (p-value < 0.05) and log |fold change| > 1.5), comparing samples with no SCNA to samples with low/high SCNA as defined by GISTIC scores. **b.** The number of differentially expressed lncRNAs per chromosome and per cancer, distinguished by color. Chromosome 1 and 17 had the most dysregulated lncRNAs associating with the greater frequency of SCNA on these chromosomes across cancers. **c.** Number of samples with structural variant breakpoints in or near (+/− 2.5kb) lncRNAs and that are also located in copy number regions, stratified by amplification or deletion status of the locus.

While SCNAs can cause the dysregulation of lncRNA expression based on gene dosage, structural variant (SV) breakpoints within a lncRNA could cause loss or gain of function^36,44^. We utilized WGS data to identify lncRNAs disrupted by SV breakpoints using a previously described combination approach involving copy number read-depth and discordant junction approach^44^ (**Online Methods**). There were 650 unique expressed lncRNA genes disrupted by SVs, 89% of which were found in only one sample (**Supplementary Fig. 4a)**. We observed 212 SV-impacted lncRNA genes located at SCNA regions (**Fig. 3c**), and 65% of lncRNAs genes disrupted by SV breakpoints in at least five samples were located at SCNA regions (**Supplementary Fig. 4b, Supplementary Table 9**). Indeed, the top-ranked SV-impacted lncRNA in both NBL and WT, *MYCNOS,* associates with the disease-driving chr2p24 amplification^46,47^ (**Supplementary Fig. 4c-d)**. In B-ALL, the SV-impacted lncRNAs: *KIAA0125* and *CDKN2B-AS1 (ANRIL)* associate with the well-studied *IGH* translocation and *CDKN2A*/*B* deletion locus (**Supplementary Fig. 4e**)^48^. The top-ranked SV-impacted lncRNA in AML, *MIR181A1HG (MONC)*, associates with a recurrent SCNA deletion on 1q and is mildly up-regulated in the AML dataset (p = 0.061, **Supplementary Fig. 4f**). *MIR181A1HG* (*MONC*) was described previously as an oncogene in acute megakaryoblastic leukemia^49,50^. Finally, we observed 30 lncRNAs with pan-cancer (n>3) expression and SV breakpoints(**Supplementary Fig. 4h**). The most number of breakpoints across unique samples was observed in *LINC00910,* which was shown previously to be essential for cell growth in the K562 cell line^51^.

### Characterization of transcriptional network perturbation mediated by dysregulated lncRNAs

To determine how lncRNAs may drive pediatric cancers, we examined the downstream impact of lncRNAs on gene regulation. We focused on identifying lncRNAs that mediate transcriptional regulation by modulating TF activity (lncRNA modulators)^52–55^. We wrote custom scripts implementing the lncMod computational framework^56^ (**Online Methods**) to first identify DE-lncRNAs and then to assess their impact on correlated expression between a TF and its target genes^21,56^ (**Fig. 4a, Online Methods**). Across all cancers studied, we identified 313,370 unique, dysregulated lncMod triplets (lncRNA-TF-target gene), representing 0.02-0.2% of possible triplets, which have significant correlation differences between a TF and target gene upon lncRNA expression dysregulation (**Supplementary Table 10-11**). This proportion was consistent with previous findings from the lncMap study in adult cancers^21^, although more triplets were identified in datasets with greater sample size (**Supplementary Table 10-11**). LncRNA modulators were categorized into one of three categories based on their impact on TF-target gene correlation; either the correlation was enhanced, attenuated, or inverted (**Fig 4a-b**). lncRNA modulators have context specific function such that for different TF-target gene pairs they could exert different types of regulation (**Supplementary Fig. 5b**). The majority of lncRNA modulators appeared to be active in only one cancer, with only 15% (138 of 923 lncRNAs) having pan-cancer activity (n>3) (**Fig. 4c**).

**Fig 4:**
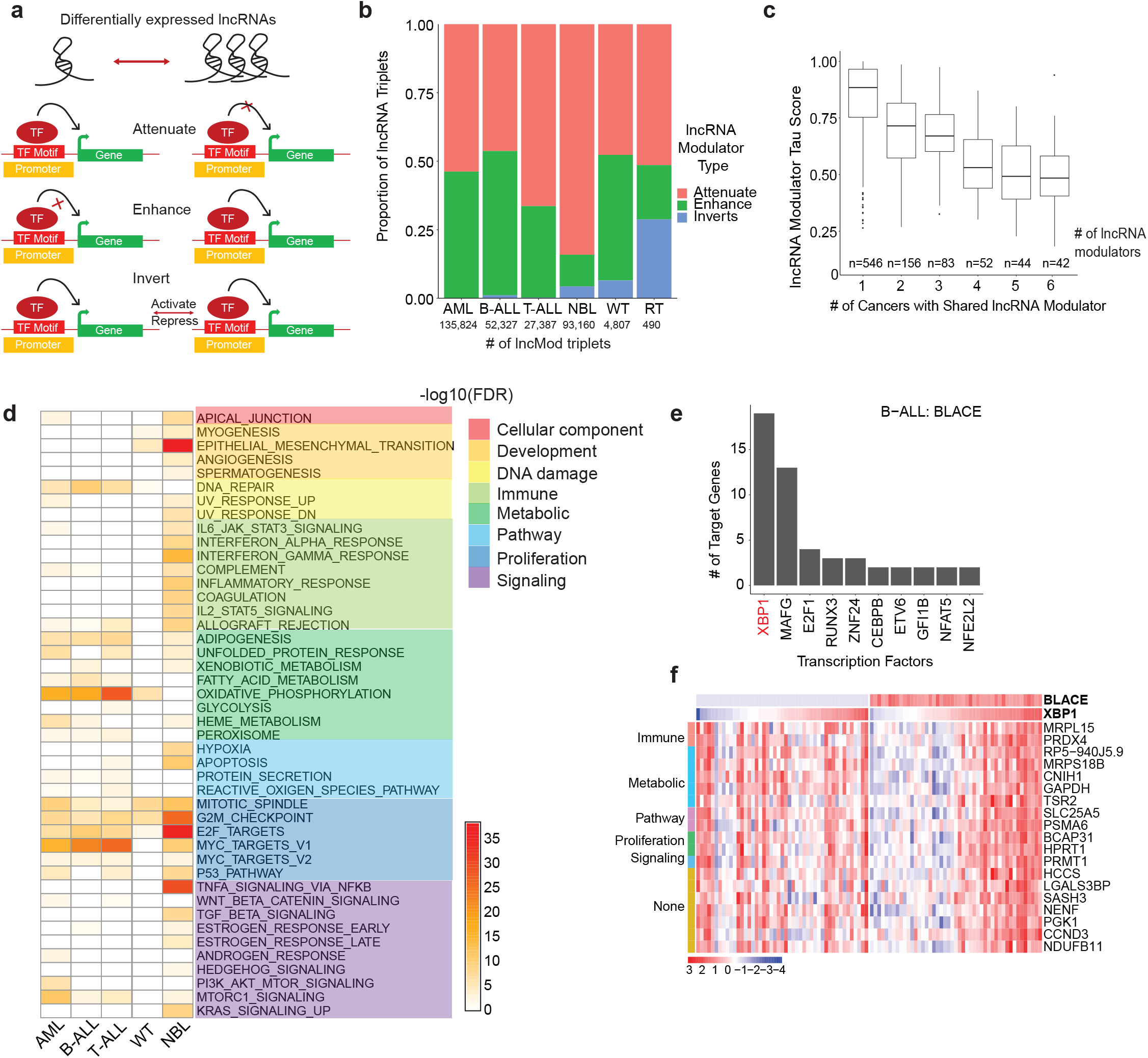
lncRNA modulators impact transcriptional networks involving proliferation. **a.** Schematic that shows the three ways (attenuate, enhance, or invert) in which differentially expressed lncRNA modulators can impact transcription factor and target gene relationships. lncRNA modulators are associated with a TF-target gene pair based on a significant difference between TF-target gene expression correlation in samples with low lncRNA expression (lowest quartile) vs samples with high lncRNA expression (highest quartile). **b.** The proportion of lncRNA modulator types associated with significantly dysregulated lncRNA modulator-TF-target gene (lncMod) triplets. The number of significantly dysregulated lncMod triplets is listed per cancer. **c.** Number of lncRNA modulators genes that are common in lncMod triplets across cancers. Common lncRNA modulator genes tend to have a lower tau score compared to lncRNA modulators only associated with one cancer. **d.** Gene set enrichment using the MSigDB Hallmark gene set, of target genes associated with lncRNA modulators in each cancer (Fisher’s exact test, FDR < 0.1). **e.** Transcription factors associated with the B-ALL expression specific lncRNA, *BLACE*, ranked based on number of regulated target genes. **f.** Expression heatmap of *BLACE* and the target genes of the XBP1 transcription factor, grouped by associated hallmark gene set, in samples within the bottom and top quartiles of *BLACE* expression in B-ALL.

To determine the biological impact of lncRNA modulators, we identified lncRNAs whose target genes were enriched in MSigDB’s Hallmark Gene Sets (HMS)^57^ (Fisher’s exact test, FDR < 0.1; **Online Methods**). Across the majority of cancers, lncRNA modulator target genes had significant enrichment in the proliferation, metabolism, and DNA damage hallmark categories (FDR range: 0.1 to 2.24×10^−36^; **Fig. 4d**). Overall, the top-enriched hallmark pathways closely mirrored those found for lncRNA modulators in adult cancers^22^. Consistent with its role in development and as an oncogene in certain cancers ^23^, the top-enriched hallmarks for the *H19* lncRNA, dysregulated in NBL, were the EMT (development) and G2M-checkpoint (proliferation) hallmarks (**Supplementary Fig. 5c**). The blood cancers exhibited strong enrichment of lncRNA modulators regulating MYC targets, which has a well-established role in leukemias^58^. Furthermore, in AML, we observed that gene targets of the myeloid-specific lncRNA, *HOTAIRM1*, were most enriched for proliferation hallmarks (**Supplementary Fig. 5d**), consistent with this lncRNA’s known role in proliferation as an oncogene in adult AML^59^.

Finally, we sought to determine potential lncRNA mechanism by identifying recurring patterns of regulation amongst lncMod triplets. To this end, we nominated candidate lncRNA-TF associations by ranking TF’s based on the number of target genes regulated by each given TF (**Supplementary Table 12**). As proof-of-concept, we were able to detect known lncRNA-TF associations such as *GAS5* with E2F4^60^ (RNA-protein), and *SNHG1* with *TP53*^61^ (RNA-RNA) amongst lncMod triplets in our study (**Supplementary Fig. 5e-f**). A notable example from the hundreds of novel associations identified is between the B-ALL specific lncRNA, *BLACE* (B-cell acute lymphoblastic leukemia expressed, tau score: 0.999) and its top associated TF, XBP1, which has known roles in pre-B-ALL cell proliferation and tumorigenesis^62^ (**Fig 4e-f)**. These predictions of lncRNA transcriptional networks provide focused avenues to elucidate the mechanisms through which lncRNAs can drive pediatric cancers.

### Defining the role of lncRNAs in childhood cancer development

Pediatric cancers arise in the context of normal human development where cells do not differentiate as they should, resulting in malignant cell transformation^63^. Some tumors are comprised of heterogenous cells that resemble varying differentiation lineages with distinct transcriptomic states due to distinct super enhancer transcription factor networks^64,65^. We sought to uncover lncRNAs associated with these varying cell lineages as they may contribute to pediatric cancer etiology. We used NBL as a model given its heterogeneity and two confirmed tumor cell states: the undifferentiated mesenchymal (MES) cells and the committed adrenergic (ADRN) cells, which can interconvert^66^. Given that NBL precursor cells, the neural crest cells, have been shown to have a more MES gene expression signature^65,66^, we hypothesized that lncRNAs correlated with an MES signature may play a role in NBL development. Using the gene set variation analysis (GSVA) method^67^ we assigned for each NBL Stage 4 sample, both a MES and ADRN score (**Online Methods**). Using hierarchical clustering (**Supplementary Figure 6a**) we categorized samples based on their primary gene expression phenotype as ADRN, MES, or mixed (**Fig 5a**). We next correlated the MES and ADRN score with lncRNA expression across NBL samples. We observed 29 lncRNAs associated with MES samples and 21 lncRNAs associated with ADRN samples (**Fig 5b**) (Spearman’s |rho| >0.6, adj. pval < 0.01). We then performed a guilt-by-association analysis^68^ to determine the potential functional pathway for these lncRNAs based on the pathway of their correlated protein coding genes (**Online Methods**). Gene set enrichment was performed using the gene ontology (GO) biological processes gene set. Intriguingly, the ADRN group of lncRNAs showed enrichment for DNA replication and cell cycle associated gene sets, whereas the MES lncRNAs were associated with organ development and immune response (**Fig 5b**). We validated these same pathway results in an independent analysis of the GMKF NBL cohort restricted to Stage 4 samples (n=67) (**Supplementary Figure 6b**). Across both TARGET and GMKF cohorts we observed 13 lncRNAs strongly associated with MES samples (**Supplementary Table 6c**), which warrant further study for their potential role in NBL development.

**Fig 5:**
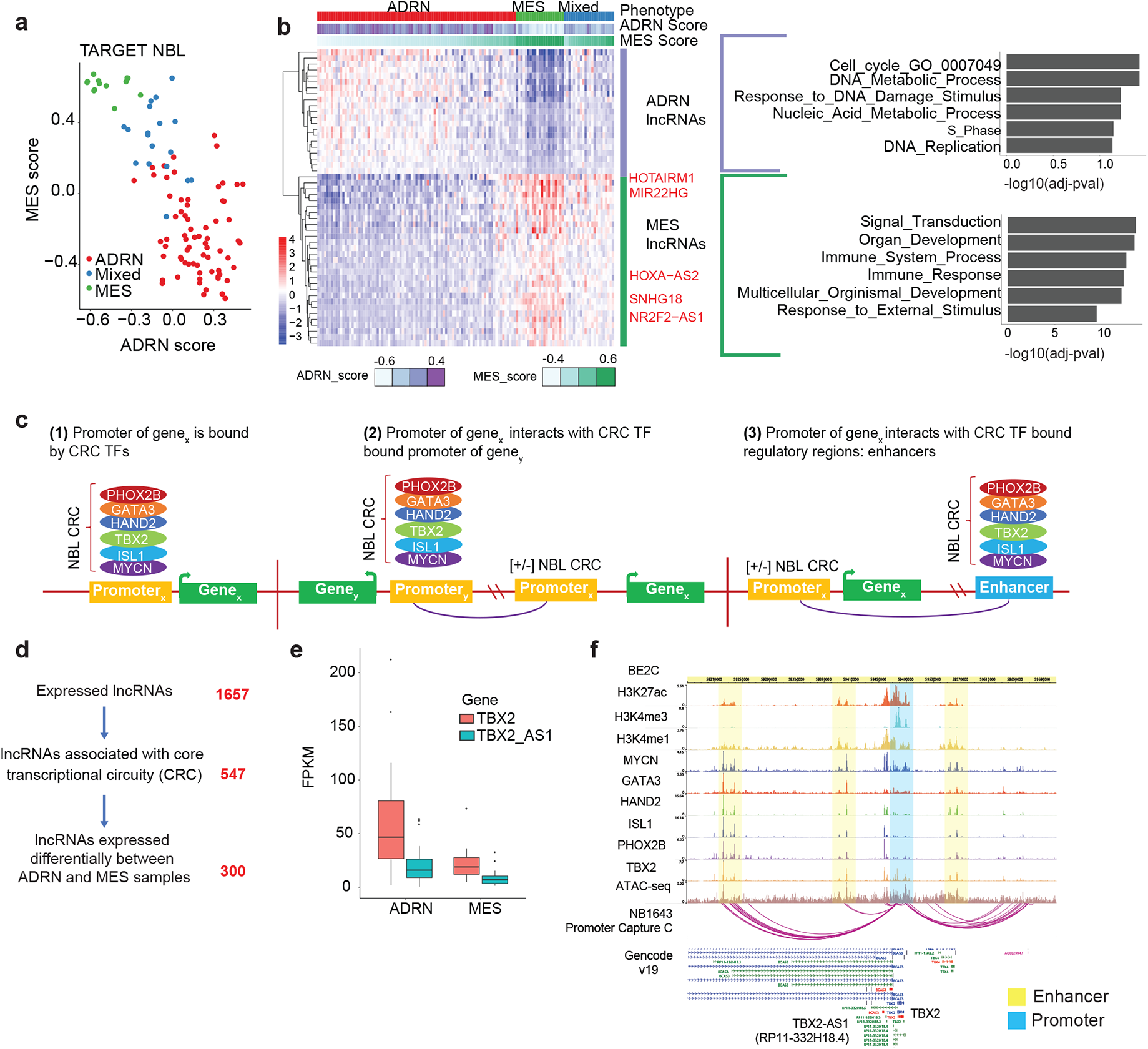
Identification of lncRNAs associated with distinct neuroblastoma cell states. **a.** The MES and ADRN signature score for TARGET NBL samples, with each sample labeled with either ADRN, Mixed, or MES phenotype based on clustering analysis. **b.** Heatmap of the expression of lncRNAs that have significant correlation with either the MES or ADRN score (|r| >0.6, pval < 0.01). lncRNAs were correlated with protein coding genes on the same chromosome and subsequent gene set enrichment analysis was performed for MES and ADRN protein coding genes separately. **c.** Schematic of how ADRN associated CRC regulated genes are identified using ChIP-seq and chromatin interaction data. We identified lncRNAs based on three types of regulation. 1) CRC transcription factors binding directly at the promoter of the lncRNA. 2) CRC TFs bind an enhancer region that interacts with a lncRNA promoter. 3) CRC TFs bind the promoter of a different gene and this promoter interacts with a lncRNA promoter. CRC TF binding was identified from ChIP-seq data, while enhancer-promoter and promoter-promoter interactions were identified from chromatin capture data. **d.** Filtering of lncRNAs expressed in NBL based on CRC TF regulation and differential expression based on sample phenotypes (ADRN or MES). **e.** Expression of *TBX2* and *TBX2-AS1* stratified by NBL sample phenotype (ADRN or MES). **f.** ChIP-seq tracks for histone marks and CRC transcription factors in the NBL cell line: BE(2)C, and promoter capture C chromatin interactions in NBL cell line: NB1643, at the *TBX2*/*TBX2-AS1* locus.

### Identification of potential cancer driver lncRNAs via integration of epigenetic data

To better identify ADRN lncRNAs we elucidated lncRNAs directly regulated by the known ADRN transcription factors (TFs): MYCN, PHOX2B, HAND2, GATA3, ISL1, and TBX2^65,69^. This set of TFs, which are co-bound and auto-regulated, are known as the core transcriptional circuitries (CRC) and drive the ADRN cell lineage in NBL^65,69^. CRC gene regulation occurs both by direct promoter binding (**Fig. 5c-1**) and by distal binding to either promoters (**Fig. 5c-2**) or enhancer regions (**Fig. 5c-3**) which then regulate the gene of interest via long-range chromatin interactions^65,69–71^. CRC-bound regulatory loci were identified from publicly available ChIP-seq data for all ADRN TFs across two MYCN-amplified NBL cell lines: SKNBE(2)C and KELLY^69,72^ (**Online Methods**). To comprehensively identify both short- and long-range CRC gene regulation, we generated high-resolution (i.e. using 4-cutter restriction enzyme DpnII) genome-wide promoter-focused Capture C^73^ in the NBL cell line NB1643. After pinpointing gene promoters interacting with CRC TF bound regulatory loci (promoters or enhancers) (**Fig. 5c, Online Methods**), we identified 547 lncRNA genes associated with the NBL CRC (**Fig 5d, Supplementary Table 13**), with only 249 of these lncRNA genes being bound by CRC TFs within their promoter regions. We further distinguished 313 ADRN lncRNAs based on differential expression (DE) between ADRN and MES samples (**Fig 5d, Supplementary Table 14**). The *TBX2-AS1* DE-lncRNA was highly correlated to the CRC TF: *TBX2* (Pearson’s r=0.77), and both are up-regulated in ADRN samples (**Fig 5e**). CRC binding is observed at both the shared promoter region of *TBX2* and *TBX2-AS1* and at an interacting distal enhancer (**Fig 5f**). TBX2 was recently shown to be involved in NBL cell proliferation,^74^ but the role of *TBX2-AS1* in NBL is unknown.

To further demonstrate the utility of this epigenetic based prioritization, we applied the same method to T-ALL, which also has a well-established set of CRC TFs (TAL1, MYB, GATA3, and RUNX1)^71^. We used available ChIP-seq and ChIA-PET data for the TAL1 mutated T-ALL cell lines, Jurkat and CCRF-CEM, to identify loci bound by the T-ALL CRC TF’s^71^ (**Online Methods**). We not only identified the known leukemia associated lncRNA PVT1, but also 9 other T-ALL CRC lncRNAs prioritized based on correlation with T-ALL PCGs and differential expression associated with a previously defined TAL1-subgroup^75^ (**Supplementary Figure 7, Supplementary Table 13-15**). Taken together, this novel data integration method nominates multiple lncRNAs with previously unknown function for further study as potential driver genes in pediatric cancer.

### Integrative multi-omic analysis prioritizes *TBX2-AS1* as a candidate functional lncRNA in NBL

To obtain a comprehensive prioritization of candidate functional lncRNAs for each cancer histotype, we integrated information for (1) tissue specific expression, (2) dysregulation due to DNA copy number aberration, and (3) regulation by CRC TFs (**Supplementary Table 16**). Here, we focus on the NBL cohort since this cancer has data available for all of the prioritization steps (**Supplementary Table 17**). The top ranked lncRNA in NBL was *MEG3*, which has a known role in both NBL and other cancers^43^. The next notable lncRNA, *TBX2-AS1,* is up-regulated due to chromosome 17q gain (**Fig 6a**), has NBL-specific expression (tau score: *TBX2*-0.807, *TBX2-AS1*-0.86; **Supplementary Fig. 8a**), and is co-regulated with *TBX2*. TBX2 has been shown to drive NBL proliferation via the *FOXM1*/*E2F1* gene regulatory network^72^ and we hypothesized that *TBX2-AS1* may play a similar role because predictions from our lncMod analysis indicated that *TBX2-AS1* impacts E2F targets and G2M checkpoint genes (**Fig. 6b**). Furthermore, the TFs primarily impacted by TBX2 knockdown^72^, MYBL2 and E2F1, were found to have the most target genes predicted to be regulated by *TBX2-AS1* (**Fig 6c-d**). Evidence for this association was further supported by the correlation (Spearman’s rho > 0.4) between *TBX2-AS1* and *TBX2*’s target TFs, including: *FOXM1*, *E2F1*, and *MYBL2* (**Supplementary Fig. 8b**). While the strong correlation between *TBX2-AS1* and *TBX2* may confound our predictions, a previous study showed positionally conserved lncRNAs^59^, such as *TBX2-AS1*, often regulate their neighboring developmental TFs (TBX2) and can play roles in genome organization and cancer^59^. Based on the promising *in silico* evidence, we prioritized *TBX2-AS1* for experimental study.

**Fig 6:**
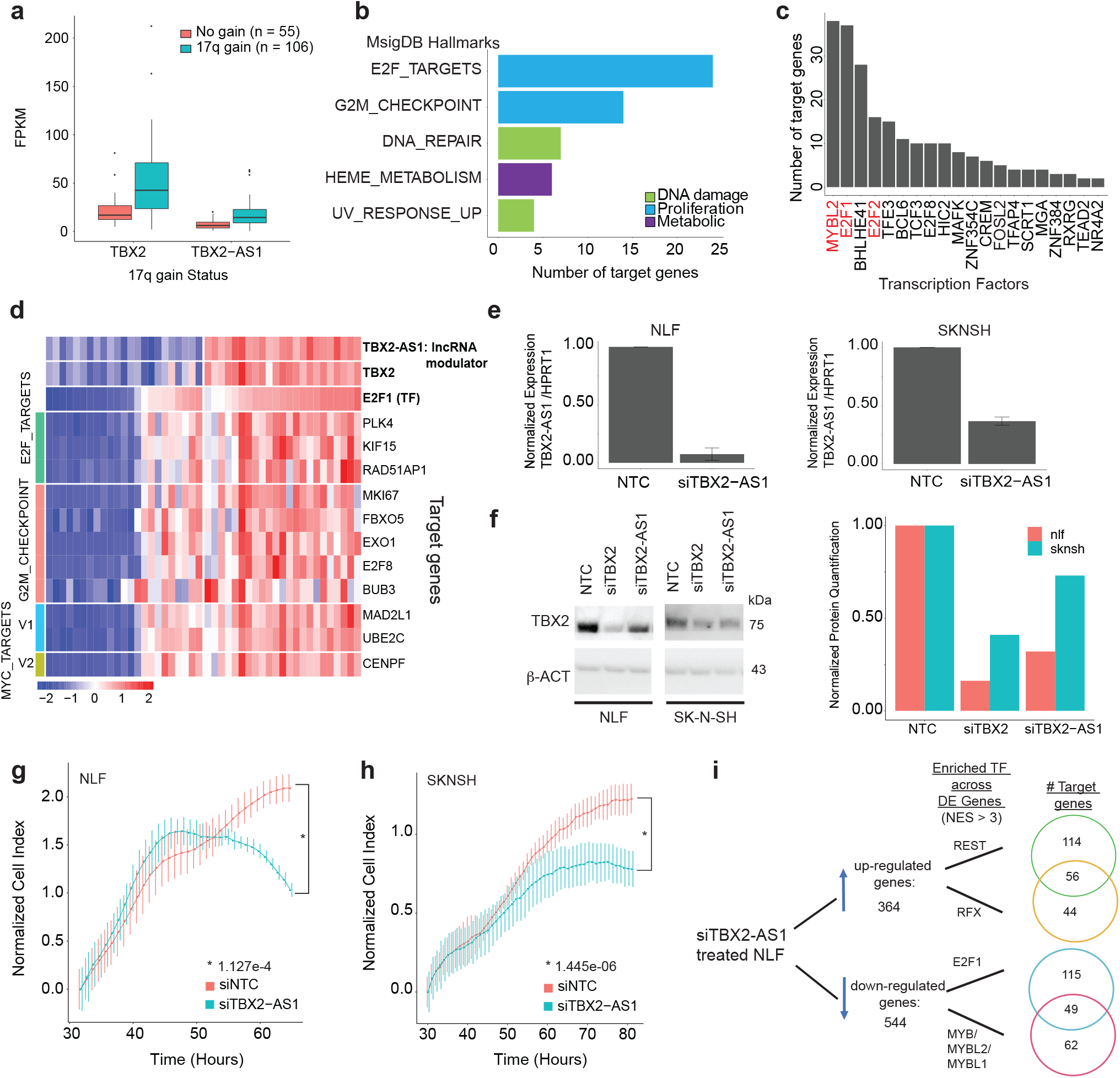
The *TBX2-AS1* lncRNA plays a role in neuroblastoma proliferation by modulating TBX2. **a.** Expression of *TBX2* and *TBX2-AS1* in NBL tumor samples with and without 17q gain. **b.** The top MSigDB Hallmarks enriched across targets genes (p-value < 0.01) regulated by *TBX2-AS1* as predicted from lncMod analysis. **c.** The transcription factors with most target genes regulated by *TBX2-AS1* as predicted from lncMod analysis. **d.** Expression of gene targets of the E2F1 transcription factor that are enriched for proliferation hallmarks, in samples with low and high *TBX2* and *TBX2-AS1* expression. *TBX2* expression is highly correlated with that of *TBX2-AS1* (Pearson’s r=0.77). **e.** siRNA knockdown efficiency of *TBX2-AS1* in the NBL cell line: NLF is 91% and in the SKNSH cell 63% knockdown was achieved. **f.** Western blot analysis of TBX2 in siTBX2 and siTBX2-AS1 treated NLF and SKNSH cell lines. **g.** Representative image of cell growth (as measured by RT-Ces assay) of the NBL cell lines: NLF and **h**. SKNSH. Cell index is normalized to time point when siRNA reagent is added at 24 hours post cell plating. **i.** Results from iRegulon analysis for genes that are up- or down-regulated upon siTBX2-AS1 treatment in NLF. Number of genes shown in Venn diagram with evidence of motif or ChIP-seq binding of the listed transcription factors.

### Silencing of *TBX2*-*AS1* inhibits cell growth of neuroblastoma cells

We assessed the role of *TBX2-AS1* using human-derived NBL cell line models. First, we evaluated *TBX2-AS1* expression across 38 NBL cell lines using RNA-seq^76^ (**Supplementary Fig 8c**). Expression of *TBX2* and *TBX2-AS1* were subsequently validated in eight cell lines using RT-qPCR (**Supplementary Fig. 8d**). We selected NLF and SKNSH models for further study based on their high *TBX2-AS1* expression and differing expression levels of *TBX2*. Silencing of *TBX2-AS1* using small interfering RNA (siRNA) achieved 92% and 63% reduction of *TBX2-AS1* expression in NLF and SKNSH, respectively (**Fig. 6e**). We also observed down-regulation of TBX2 protein levels in the siTBX2-AS1 treated cells for both cell lines (**Fig. 6f**). Given the known role of TBX2 in NBL cell proliferation^74^, we measured cell growth of siTBX2-AS1 treated NBL cells to determine if *TBX2-AS1* has similar function. When the non-targeting control (siNTC) treated cells reached confluence, the siTBX2-AS1 treated cell index was reduced by 42.6% and 36.8% (n=3, p < 0.01) in the NLF and SKNSH cell line, respectively (**Fig. 6g-h, Supplementary Fig. 8e**). Live cell imaging using the IncuCyte revealed changes in cell morphology for siTBX2-AS1 treated NLF cells, featuring an appearance of disrupted cell to cell adhesion and elongated cell body (**Supplementary Fig. 8f**). To identify pathways impacted by *TBX2-AS1* knockdown, we performed total RNA sequencing in triplicate of NLF cells and compared gene expression in control (siNTC) vs siTBX2-AS1 treated cells (**Supplementary Fig. 8g**). Gene set enrichment analysis (GSEA) of the 364 significantly up-regulated genes (log-fold change > 1.5, adj pval < 0.1) revealed enrichment (FDR < 0.1) for hallmarks associated with inflammation including: TNFA signaling and interferon gamma response (**Supplementary Table 18**). Across the 544 down-regulated genes, E2F target genes hallmark was most enriched. To determine whether differentially expressed genes shared common regulation, we used the iRegulon program^77^, to search for TF motifs and ENCODE ChIP-seq tracks upstream of genes (**Online Methods**). Using a normalized enrichment score (NES) of at least 3, we observed motif enrichment for the neuronal differentiation repressor REST and the RFX family of transcription factors in 59% of siTBX2-AS1 up-regulated genes (**Fig. 6i**). In 42% of downregulated genes, the top enriched TFs were MYBL2 and E2F1, corroborating our GSEA results. Moreover, both the growth assays and gene expression profiling confirmed our lncMod results, which showed that *TBX2-AS1* impacts NBL proliferation by modulating target genes of E2F1 and MYBL2 (**Fig. 6b-d**). These data thus demonstrate the utility of our integrative lncRNA characterization and prioritization approach for future validation experiments across all cancers considered in this study. Furthermore, we uncovered a functional role for *TBX2-AS1* in NBL proliferation likely mediated via the regulation of TBX2 and its known target genes: E2F, MYBL2, and REST^72^.

## Discussion

LncRNAs have emerged as important regulators of gene expression and their dysregulation can impact key cancer pathways and drive tumorigenesis^1–4^. Despite this, relatively few lncRNAs have been experimentally characterized and the landscape of lncRNA expression across pediatric cancers has been previously unknown. In this study, we explored lncRNA expression, cancer association, and regulatory networks across 1,044 pediatric leukemias and solid tumors, representing six different cancer types. The breadth of samples and cancer types included allowed for robust identification of novel, cancer-specific, and developmental lncRNAs. Furthermore, we used systems modelling to identify expression patterns for both up- and downstream lncRNA gene regulation. Altogether we provide multi-dimensional insight into the predicted biological and functional relevance of lncRNAs by integrating WGS, ChIP-seq, chromatin capture, and predictions of transcriptional networks.

Analysis of the lncRNA landscape across pediatric cancers revealed the histotype and context-specific nature of lncRNAs. We report a total of 2,657 robustly expressed lncRNAs across the six cancer types studied. This number is notably smaller than reports from pan-cancer studies of adult malignancies^15,17^, likely due to the smaller number of cancer types studied here and conservative expression threshold applied. However, similar to our findings in adult cancers, 43% (1,142/ 2,657) of expressed lncRNAs exhibited tissue-specific (TS) expression across pediatric cancers. Indeed, lncRNAs had significantly greater tissue specificity than protein coding genes, making them more ideal candidates as biomarkers. Currently there is one lncRNA, *PCA3*, that is FDA-approved as a biomarker for prostate cancer^78^ and multiple trials investigating ncRNAs in cancer prognostics are underway^79^. In this study, the top five most TS lncRNAs per cancer were sufficient to differentiate each cancer histotype. Furthermore, we identify lncRNAs specific to distinct cell lineages within NBL, suggesting there is potential for lncRNAs to be used as highly sensitive markers to differentiate cancer subtypes more accurately.

Typically, investigation of lncRNA dysregulation involves comparing lncRNA expression between cancer and normal control samples and is an analysis that amply yields adult-cancer associated lncRNAs^15^. However, the lack of normal expression controls for the majority of pediatric cancers^36^ is a major complication in defining pediatric cancer-associated lncRNAs. To overcome this, we leveraged information about how pediatric cancers are epigenetically regulated. In particular, NBL, is composed of two cells lineages representing different development stages and each with distinct super-enhancer transcription factor networks. Given the tie between organogenesis and tumorigenesis in pediatric cancer^63^, we hypothesized that lncRNAs associated with these cell states may also be involved in NBL development. After correlation and pathway analysis, we discovered that lncRNAs associated with the mesenchymal cell lineage had enrichment for organogenesis gene sets, while adrenergic-associated lncRNAs were predicted to be involved in proliferation based on enrichment for DNA replication and cell cycle gene sets. The majority of NBL samples have cells with an adrenergic gene expression signature, which could suggest that ADRN lncRNAs are major drivers of disease and thus potential therapeutic targets. To better identify these ADRN lncRNAs, we integrated ChIP-sequencing of core regulatory (CRC) transcription factors for ADRN cells with our expression data to identify cancer driver lncRNAs. CRC TFs bind to cell-type-specific enhancers and regulate the expression of cell-type-specific genes^80^. By taking advantage of this information we were able to prioritize lncRNAs likely to be important for cancer cell identity based on CRC TF regulation. CRC TFs have been well defined for NBL and T-ALL^69,71^; however the fact that they largely bind enhancer regions necessitated that we also use chromatin interaction data to accurately determine regulated genes. Incorporation of these datasets allowed us to identify 2-fold more CRC regulated lncRNAs in NBL and 3-fold in T-ALL as compared to using just ChIP-seq data alone, which restricts lncRNA identification to those with CRC TFs bound at their promoter. Notably, there were ten common CRC-regulated lncRNAs between NBL and T-ALL, and an important next step for further identification of pan-pediatric cancer associated lncRNAs is application of this novel analysis to a broader set of pediatric cancers.

While upstream regulation can help nominate cancer-associated lncRNAs, determining the mechanism through which dysregulated lncRNAs impact downstream target genes is also crucial. However, prediction of lncRNA function is limited given that very few lncRNA mechanisms have been fully established and lncRNAs lack conserved sequence and structure^81^. Many studies instead use correlated protein coding gene expression as a proxy to define lncRNA pathways, but this approach often results in many false positives and does not provide mechanistic insight^81^. To address this, we used the lncMod method^21,56^ to model the functional mechanism of dysregulated lncRNAs by examining correlated changes in transcription factor to target gene regulation. We used motif presence and regression analysis to identify TF-target gene relationships, though future studies will be strengthened by incorporating TF ChIP-seq data, when it becomes more widely available for pediatric cancers. Nevertheless, we were able to successfully associate lncRNAs to TFs with known interactions, such as SNHG1 with TP53, while also providing a prioritized list of novel associations that serve as a starting point for future experimental studies such as RIP/MS^82^ and ChiRP-seq^83^. Finally, while our lncMod analysis was focused on transcriptional regulation, the addition of microRNA binding and RNA-binding protein data, as utilized in adult cancers^22^, is an important next step in understanding how lncRNAs impact post-transcriptional regulation in pediatric cancers.

Our study delineated high confidence lncRNA expression across pediatric cancers within the restrictions set by the sequencing depth and RNA-seq type available per cancer dataset. We required RNA-seq samples included in our study to have at least 10 million reads and read length of at least 75 bp; and with the exception of the T-ALL samples, all samples were poly-A selected. Future studies involving total RNA-seq, greater sequencing depth, and longer read sizes could capture a larger diversity and more accurate set of expressed lncRNAs by accounting for non-polyadenylated genes and identifying scarcer or temporally expressed lncRNAs. Nevertheless, our high confidence set of lncRNAs are very likely to be functional given that low or rare expression can be an indicator of transcriptional noise^84^. In addition to having a limited number of RNA matched WGS samples, the Complete Genomics short read technology limits the detection of structural variants based on size as previously described^36,44^. The use of long-read sequencing and greater sequencing depth in future studies will enable more accurate copy number and structure variant detection in pediatric cancers.

Finally, multi-dimensional integration of our computational predictions resulted in the nomination of functionally relevant lncRNAs in each pediatric cancer. We annotated tissue specificity, copy number, pathway, and likely targets for these lncRNAs, providing a solid foundation for mechanistic studies. As proof-of-principle, we demonstrate that the top-prioritized tissue-specific and copy number dysregulated lncRNA, *TBX2-AS1,* impacts NBL cell growth, validating our approach, while transcriptomic profiling corroborated our pathway predictions. Knockdown of *TBX2-AS1*, showed downregulation of genes regulated by E2F1 and MYBL2, the same TFs impacted upon TBX2 knockdown^72^. Future studies could reveal whether *TBX2-AS1* modulates TBX2 through direct binding or by impacting transcriptional regulation at their shared locus. *TBX2-AS1* was previously shown to be among a group of lncRNAs which are positionally conserved and near developmental associated TFs^59^. This group of lncRNAs and their neighboring TFs, typically have tissue specific expression, can be involved in cancer development, and affect each other’s expression^59^, all of which we observed for TBX2 and *TBX2-AS1*. Together these genes contribute to the proliferative state of NBL cells and could have potential as novel therapeutic targets.

Altogether, this study provides a comprehensive characterization of lncRNAs across pediatric cancers and serves as a rich resource for future mechanistic studies; these data may aid in the selection of cancer biomarkers and candidate therapeutic lncRNA targets.

## Online Methods

### RNA-seq data processing

A comprehensive RNA-seq analysis pipeline was used on all samples (Supplementary Table 1, Supplementary Fig 1). First FASTQC was run on all samples and any samples that had a Phred score < 30 for more than 25% of read bases were removed. Samples were then aligned using STAR_2.4.2a ^85^ with the following parameters: “STAR --runMode alignReads --runThreadN 10 -- twopassMode Basic --twopass1readsN -1 --chimSegmentMin 15 --chimOutType WithinBAM –genomeDir X--genomeFastaFiles ucsc.hg19.fa --readFilesIn fasta1 fasta2 --readFilesCommand zcat --outSAMtype BAM SortedByCoordinate --outFileNamePrefix X --outSAMstrandField intronMotif --quantMode TranscriptomeSAM GeneCounts --sjdbGTFfile gencode.v19.annotation.gtf --sjdbOverhang X.” To assess the quality of the aligned RNA-seq data we ran MultiQC ^86^, and removed samples with < 70% uniquely mapped reads and < 10 million mapped reads.

### Gene/transcript mapping and quantification

To map reads to genes and quantify gene expression we ran StringTie 1.3.3 ^37^. StringTie involves three steps, first quantifying expression of both known and novel gene transcripts using an annotation guided approach. We used the Gencode v19 gene annotation to guide gene detection.1) “stringtie bamfile -G gencode.v19.annotation_stringtie.gtf -B --rf -o out.gtf -A gene_abund.tab -C cov_refs.gtf -p 10.” In the second step, StringTie merges the gene annotation across all samples such that there is a uniform annotation for known and novel gene transcripts in one transcriptome gtf file. 2) “stringtie All_PanTARGET_PreMerge_StringTie_Files.txt --merge -G gencode.v19.annotation_stringtie.gtf -o StringTie_PanCancer_AllMergedTranscripts.gtf.” Finally, StringTie is run again to quantify expression using the PanTarget transcriptome gtf file and de novo gene transcript detection is turned off. 3) “stringtie bamfile -G StringTie_PanCancer_AllMergedTranscripts.gtf -B -e --rf -o out.gtf -A gene_abund.tab -C cov_refs.gtf -p 10”

### Comparison of pan-TARGET transcriptome with reference annotation

Novel transcripts were assigned as an isoform of a known gene based on exonic overlap (>50% by bp) with genes in either the GENCODE v19 or RefSeq v74 databases using custom Python scripts. Any remaining novel transcripts were assigned as novel genes (MSTRG_Merged.# or MSTRG.#) based on overlapping exon positions. Novel genes were further filtered based on read coverage, in that we required that at least one transcript for a novel gene have more than one exon with at least 5 reads in at least 20% of samples per cancer. High confidence novel genes were required to have at least 3 exons. Finally, for all transcripts (known and novel), to obtain gene level quantification, transcript FPKM and count values were summed to get a gene level value.

### Prediction of novel gene coding potential and lncRNA gene annotation

We predicted coding potential of novel transcripts using the PLEK v1 algorithm tool ^39^. PLEK uses a support vector machine (SVM) for a binary classification model to distinguish a lncRNA versus a coding mRNA. The features used as input for the SVM are calibrated k-mer usage frequencies of a transcript’s sequence. PLEK has previously been validated on RefSeq mRNAs and GENCODE lncRNAs (the main reference annotations used in our study) and has achieved >90% accuracy in predicting gene coding potential ^39^. To further delineate lncRNAs, we removed any predicted novel non-coding transcripts that were < 200bp (sum of total exon length). We updated the gene type of GENCODE v19 genes with the gene type of genes that had matching gene names in GENCODE v29. Additionally we filtered out lncRNA genes that have been deprecated in Gencode v29. Finally, some lncRNA genes in Gencode v19, have both a lncRNA and small RNA transcript. For these 147 cases we did not include the small RNA transcript when summing gene transcripts to obtain gene level expression.

### Tissue specific gene expression

The tau score, a measure of the tissue specific expression of a gene was calculated as described by Yanai et. al^41^. The formula for the score is listed below. x_i_ is defined as the mean expression of a gene in a particular cancer and n is the total number of cancers considered, in this case n = 6.

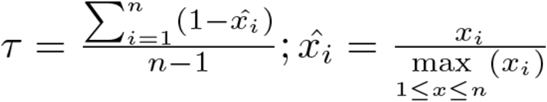

### CNV detection, processing, and impact on gene expression

Copy number calls were made by Complete Genomics (CGI) from WGS for NBL, WT, AML, and B-ALL. We used CGI files“somaticCnvDetailsDiploidBeta” containing ploidy estimates and tumor/blood coverage along 2kb bins across the genome. To create segmentation files, we used custom scripts to reformat CGI coverage data to meet requirements of the “copynumber” R bioconductor package as previously described^44^. We used the winsorize function in this package, which performs data smoothing and segmentation via a piecewise constant segmentation (pcf) algorithm (kmin =2 and gamma= 1000). Segmentation files were visualized using the R package svpluscnv (https://github.com/ccbiolab/svpluscnv) https://doi.org/10.1093/bioinformatics/btaa878. We then ran GISTIC2.0, using segmentation data as inputs and parameters: “GISTIC2 -v 30 -refgene hg19 -genegistic 1 -smallmem 1 -broad 1 -twoside 1 - brlen 0.98 -conf 0.90 -armpeel 1 -savegene 1 -gcm extreme -js 2 -rx 0”. To determine which genes copy number impacts, we intersected CNV regions listed in the “all_lesions.conf_90.txt” file from GISTIC output with gene positions. We used section 1 from the “all_lesions.conf_90.txt” file to assign a binary descriptor to each gene as either being not amplified or deleted (CNV-no) if the sample had actual copy gain 0 for the region containing the gene. We assigned CNV-yes if the region containing the gene was amplified or deleted, which included samples with actual copy gain 1 or 2, where 1 indicates low level copy number aberration (exceeds low threshold of copy number: 1: 0.1<t< 0.9) and 2 indicates a high level of copy number aberration, CNV exceeds high threshold (t>0.9) according to GISTIC. To determine CNV impact on gene expression, we assessed differential expression of the gene in samples from the two groups (CNV yes or no) using Wilcoxon rank sum test (p < 0.01). Genes were considered to have evidence of differential expression due to copy number if the absolute value of the log2 fold change between the two groups was > 0.58 and p < 0.05.

### Structural variant detection and filtering

Structural variants were identified from WGS as previously described^44^. Somatic sequence junctions that were completely absent in the normal genome are reported by Complete Genomics (CGI) in the somaticAllJunctionsBeta file. To obtain a high confidence set of junctions, where there is a likely true physical connection between the left and right sections of a junction, the following filtering was applied by CGI to obtain the highConfidenceSomaticAllJunctionsBeta.

1. DiscordantMatePairAlignments ≥ 10 (10 or more discordant mate pairs in cluster
2. JunctionSequenceResolve = Y (local de novo assembly is successful)
3. Exclude interchromosomal junction if present in any genomes in baseline samples (FrequencyInBaseline > 0)
4. Exclude the junction if overlap with known underrepresented repeats (KnownUnderrepresentedRepeat = Y): ALR/Alpha, GAATGn, HSATII, LSU_rRNA_Hsa, and RSU_rRNA_Hsa
5. Exclude the junction if the length of either of the side sections is less than 70 base pairs.

Further filtering of these high confidence structural variants included removing rare/common germline variants that passed the CGI filters. We used the Database of Genomic Variants (DGV v. 2016-05-15, GRCh37) in order to remove SVs that had at least 50% reciprocal overlap with DGV annotated common events and were type matched.

### Structural variant analysis

To obtain a comprehensive landscape of SVs we combined both the sequence junction and copy number read depth approaches to identify SVs, with co-localizing break points being orthogonally validated. Recurrence of SVs was considered based on overlap with genes from our pan-pediatric cancer transcriptome. Genomic overlap between SVs and genes was determined using the bedtools intersect tool (default parameters). Variants were assigned to genes based on if the sequence junction (left/right position) + 100 bp overlapped gene coordinates +/− 2.5kb. Genes were then ranked based on the number of unique samples per cancer with a SV breakpoint.

### Gene signature analysis

We obtained a list of genes associated with the mesenchymal (MES) and adrenergic (ADRN) NBL cell types from GEO (GSE90805). We then used the GSVA R package^67^ with the Poisson kernel (kcdf) parameter to assign a score per sample representing the total expression enrichment of genes associated with either the MES or ADRN cell types. We performed hierarchical clustering to divide NBL samples into three groups (MES, ADRN or mixed phenotype) based on expression of MES and ADRN genes using the pheatmap R package and cutting the dendrogram at n=3. We correlated the MES and ADRN score with lncRNA expression across Stage 4 NBL TARGET cohort and GMKF cohort samples separately and identified lncRNAs as having significant correlation based on absolute value Spearman’s rho > 0.6. These lncRNAs were then labeled as MES or ADRN based on significant correlation with either the MES or ADRN score. We next repeated score correlation with PCGs. We performed a guilt-by-association analysis assigning MES/ADRN PCGs and by association their correlated MES/ADRN lncRNAs (Spearman rho > 0.5) to pathways using Fisher exact test, FDR < 0.1 for gene sets in the gene ontology (GO) biological processes collection.

### ChIP-seq data analysis

To determine which lncRNAs are regulated by transcription factors involved in the core regulatory circuitry (CRC) we utilized previously generated and analyzed histone and transcription factor ChIP-sequencing data for NBL and T-ALL. For NBL, we used peak files for our previously generated histone ChIP-seq data of: H3K27ac, H3K4me1, H3K4me3 for the BE(2)C cell line^87^, available on GEO: GSE138315. We downloaded raw sequencing files for CRC transcription factor ChIP-seq data for MYCN, PHOX2B, HAND2, GATA3, TBX2, and ISL1 for the BE(2)C and KELLY cell lines from GEO: GSE94822^69^ and selected peaks with q-value < 0.001 for further analysis. We identified regions in the genome where at least 4/6 of the transcription factors overlapped. This was obtained using the homer mergePeaks tool: “mergePeaks -d 1000 -cobound 6 bed_file1… bed_file6” and the resulting coBoundBy4 output file. For the T-ALL CRC we obtained overlapping CRC transcription factor loci for TAL1, GATA3, and RUNX1 from the study by Sanda et. al^71^, GEO: GSE29181 for both the Jurkat and CCRF-CEM cell lines and integrated ChIP-seq data for the MYB transcription factor from GEO: GSE59657^70^, only available in the Jurkat line. We selected loci for further analysis if they were bound by TAL1, GATA3, and RUNX1 as previously annotated by Sanda et. al.

### Identification of CRC transcription factor regulated genes

To identify genes regulated by the NBL or T-ALL CRC we considered CRC TF binding at both the gene’s promoter and other regulatory region interacting with the gene’s promoter. We first overlapped CRC regions using bedtools intersect with gene transcript promoter regions, which we defined as 3000bp upstream and downstream of the transcripts first exon. For NBL, we then utilized the promoter-focused Capture C data, inclusive of all interactions within 1Mb on the same chromosome, to identify genomic regions that were both bound by NBL CRC TFs and interacting with a gene’s promoter. To determine this, we used bedtools intersect to determine overlap (minimum 1bp) between CRC bound loci with loci involved in chromatin interactions. From these regions, we determined which interacting regions corresponded with a lncRNA promoter region. We performed a similar analysis in T-ALL, however we utilized publicly available SMC1 (cohesin) ChIA-PET data available on the ENCODE project to consider chromatin interactions.

### Promoter-focused Capture C data generation

High resolution promoter-focused Capture C was performed in the neuroblastoma cell line, NB1643, (untreated) in triplicate. Cell fixation, 3C library generation, capture C, and sequencing was performed as described by Chesi et. al (2019) and Su et al (2020). For each replicate, 10^7^ fixed cells were centrifuged to cell pellets and split to 6 tubes for a pre-digestion incubation with 0.3%SDS, 1x NEB DpnII restriction buffer, and dH2O for 1hr at 37°C shaking at 1,000rpm. A 1.7% solution of Triton X-100 was added to each tube and shaking was continued for another hour.10 ul of DpnII (NEB, 50 U/µL) was added to each sample tube and continued shaking for 2 days. 100uL Digestion reaction was then removed and set aside for digestion efficiency QC.The remaining samples were heat inactivated incubated at 1000 rpm in a MultiTherm for 20 min, at 65°C to inactivate the DpnII, and cooled on ice for 20 additional minutes. Digested samples were ligated with 8 uL of T4 DNA ligase (HC ThermoFisher, 30 U/µL) and 1X ligase buffer at 1,000 rpm overnight at 16°C. The ligated samples were then de-crosslinked overnight at 65°C with Proteinase K (20 mg/mL, Denville Scientific) along with pre-digestion and digestion control. Both controls and ligated samples were incubated for 30 min at 37°C with RNase A (Millipore), followed by phenol/chloroform extraction, ethanol precipitation at -20°C, then the 3C libraries were centrifuged at 3000 rpm for 45 min at 4°C to pellet the samples. The pellets of 3C libraries and controls were resuspended in 300uL and 20μL dH2O, respectively, and stored at −20°C. Sample concentrations were measured by Qubit. Digestion and ligation efficiencies were assessed by gel electrophoresis on a 0.9% agarose gel and also by quantitative PCR (SYBR green, Thermo Fisher).

Isolated DNA from 3C libraries was quantified using a Qubit fluorometer (Life technologies), and 10 μg of each library was sheared in dH2O using a QSonica Q800R to an average fragment size of 350bp.QSonica settings used were 60% amplitude, 30s on, 30s off, 2 min intervals, for a total of 5 intervals at 4 °C. After shearing, DNA was purified using AMPureXP beads (Agencourt). DNA size was assessed on a Bioanalyzer 2100 using a DNA 1000 Chip (Agilent) and DNA concentration was checked via Qubit. SureSelect XT library prep kits (Agilent) were used to repair DNA ends and for adaptor ligation following the manufacturer protocol. Excess adaptors were removed using AMPureXP beads. Size and concentration were checked again by Bioanalyzer 2100 using a DNA 1000 Chip and by Qubit fluorometer before hybridization. One microgram of adaptor-ligated library was used as input for the SureSelect XT capture kit using manufacturer protocol and custom-designed 41K promoter Capture-C probe set. The quantity and quality of the captured libraries were assessed by Bioanalyzer using a high sensitivity DNA Chip and by Qubit fluorometer. SureSelect XT libraries were then paired-end sequenced on Illumina NovaSeq 6000 platform (51bp read length) at the Center for Spatial and Functional Genomics at CHOP.

### Promoter-focused Capture C data analysis

Paired-end reads from each replicated were pre-processed using the HICUP pipeline (v0.5.9), with bowtie2 as aligner and hg19 as the reference genome. The unique ditags output from HiCUP were further processed by the chicagoTools bam2chicago.sh script before significant promoter interaction calling. Significant promoter interactions at 1-DpnII fragment resolution were called using CHiCAGO (v1.1.8) with default parameters except for binsize set to 2500. Significant interactions at 4-DpnII fragment resolution were also called using CHiCAGO with artificial baitmap and rmap files in which DpnII fragments were concatenated *in silico* into 4 consecutive fragments using default parameters except for removeAdjacent set to False. Interactions with a CHiCAGO score > 5 in either 1-fragment or 4-fragment resolution were considered as significant interactions. The significant interactions were finally converted to ibed format in which each line represents a physical interaction between fragments.

### Differential gene expression analysis for T-ALL subtypes

We identified differentially expressed genes using the DESeq2 tool. We compared expression between the TAL1 subgroup and non-TAL1 subgroup, defined by Liu, et al^75^. We ran DESeq2 using default parameters and considered genes as significantly differentially expressed if their absolute value of the log2 fold change was > 0.58 and their Benjamini-Hoschberg adjusted-p value was < 0.01.

### lncMod implementation: transcription factor target gene regulation

We developed custom Python scripts to implement the general framework of the lncMod method. The first part of this framework involved determining transcription factor target gene regulation specific to each cancer. Target genes here are defined as any protein coding or lncRNA gene and excludes pseduogenes and small RNAs. Given that ChIP-seq binding profiles for the majority of transcription factors were not available for tissues associated with each of these cancers we instead used transcription factor motif analysis as a proxy. We utilized motifs in the JASPAR database^88^ and predictions of binding across the genome determined by FIMO and available in the UCSC genome database: http://expdata.cmmt.ubc.ca/JASPAR/downloads/UCSC_tracks/2018/hg19/tsv/. For each transcript we determined potential regulatory transcription factors based on the presence of predicted binding motifs in the gene promoter region. Promoter regions were defined as regions 3000 bp upstream and downstream of the transcript’s first exon. Next we selected transcription factors based on their expression in each cancer and then performed linear regression considering the expression of the transcription factor and target gene specific to each cancer. We adjusted the false discovery rate due to multiple testing using the Benjamini-Hochberg method and selected TF-target gene pairs with significantly associated expression (adjusted p-value < 1e-5).

### Identification of lncRNA modulators

To identify transcriptional perturbation, we first delineated genes (TF, target genes, or lncRNAs) that had high expression variance (IQR > 1.5). For each differentially expressed lncRNA in each cancer we calculated the following, as has been done in previous studies^21,22,56^. For a given cancer and given lncRNA we sorted samples in the cancer based on the given lncRNAs expression (low to high). We then determined the correlation (Spearman’s rho) between the expression of all transcription factor and target gene pairs previously identified in the given cancer. This correlation was calculated for the 25% of samples with the lowest lncRNA expression and separately for the 25% of samples with the highest expression for the given lncRNA. To ensure that we observed TF-target gene regulation we required that the correlation between the TF-target pair in either the low or high lncRNA expressing group was at least R>0.4. We only further evaluated the lncRNA TF-target gene triplet if the correlation difference between the low and high lncRNA expression group was R>0.45. To formally compare the correlation difference we first normalized the correlation using the Fisher r to z transformation. Then we calculated the rewiring score, z-statistic, as previously described ^21^, which is used to describe the degree of regulation change between the TF and target gene.

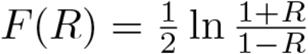

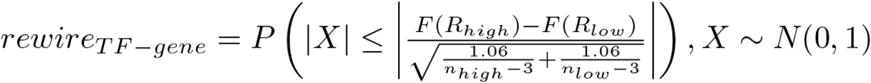

As a departure from what is described by Li et. al (lncMod method)^56^, we used permutation analysis to robustly assess the significance of the rewire score in the context of multiple hypothesis testing as described by Sham et. al^89,90^. We randomly shuffled target gene expression (TF-target gene pair labels) and calculated the rewire score P value across all TF-target gene pairs per given lncRNA. We kept the smallest observed P value and repeated the permutation 100 times. This empirical frequency distribution of the smallest P values was then compared to the P value in our real data to calculate an empirical adjusted P value (adj P value) as given by the formula below, where r is the number of permutations where the smallest P value are less than our actual P value and n is the number of permutations.

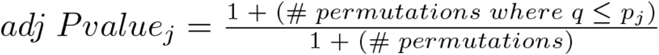

The lncRNA-TF-target gene triplets, with adjusted p < 0.1 were considered significant. Datasets with smaller sample sizes had lower statistical power and thus fewer significant triplets. Triplets were then classified into three patterns based on correlation changes between the low and high expressing lncRNA group: increased correlation – enhanced, decreased correlation – attenuated, and inverted – positive to negative correlation and vice versa. We annotated lncRNA target genes as cancer genes based on if they were listed in the COSMIC database or a complied list from Chiu et. al^22^.

### Cell lines and reagents

NBL cell lines were obtained from the American Type Tissue Culture Collection (ATCC) and grown in RPM1-1640 with HEPES, L-glutamine and phenol red, supplemented with 10% FBS, 1% L-glutamine in an incubator at 37°C with 5% CO_2_. Cell line identity was confirmed biennially through genotyping and confirmation of STR (short tandem repeat) profiles, while routine testing for Mycoplasma contamination was confirmed to be negative.

### siRNA and growth assays

The NBL cell lines, NLF and SKNSH, were plated in a 96-well RTCES microelectronic sensor array (ACEA Biosciences, San Diego, CA, USA). Cell density measurements were made every hour and were normalized to 24 hours post-plating (at transfection time). We used siRNAs to knockdown the expression of genes in NLF and SKNSH. The siRNAs utilized were either a non-targeting negative control siRNA (Silencer^TM^ Select Negative Control siRNA, cat #4390843), TBX2-AS1 Silencer^TM^ Select siRNA (cat # n514841), and SMARTpool: ON-TARGETplus PLK1 siRNA (cat # L-003290-00-0010). Transfection of cells was done using the DharmaFECT 1 transfection reagent (cat # T-2001-02). siRNA at a concentration of 50nM and 2% (NLF) and 4% (SKNSH) DharmaFECT was added to RPMI medium without 10% FBS or any antibiotic separately and then incubated at room temperature for 5 minutes. The siRNA medium was then added to the DharmaFECT and incubated for another 20 minutes to form a complex. This solution was then mixed with our normal growth media and applied to cells 24 hours after they had been initially plated. All experiments were repeated in triplicate, with technical replicates (n=3) being averaged per biological replicate.

### Real time quantitative PCR

Total RNA was extracted from NBL cells using miRNeasy kit (Qiagen) and the provided protocol for animal cells. The concentration of RNA was determined with the Nanodrop (Thermo Scientific). cDNA synthesis was performed using the SuperScript^TM^ First-Strand Synthesis System for RT-PCR using the SuperScript^TM^ reverse transcriptase (Invitrogen). 5-20ng of cDNA were mixed with the TaqMan Universal PCR Master Mix (Thermo Fisher Scientific) and TaqMan probes/primers for either TBX2-AS1 (Hs00417285_m1) or the house keeping gene, HPRT1 (Hs02800695_m1). Gene expression from these reactions were measured using RT-qPCR and TBX2-AS1 expression was normalized to HPRT1 expression.

### NLF gene knockdown expression profiling

Total RNA was isolated from the NLF cell line 48 hours post treatment with siTBX2-AS1 and non-targeting control samples, siNTC, (three biological replicates per condition) and 1000 ng/sample was used as input for library preparation with the TruSeq Stranded mRNA Sample Prep Kit from Illumina (with Ribo-Zero treatment). RNA-seq libraries were sequenced on the Nextseq 500 at depth 10 million reads per sample minimum. Library prep and sequencing was performed by the Sidney Kimmel Cancer Center Genomics Facility of Thomas Jefferson University. Sample and read quality was assessed using FastQC and reads were aligned and mapped using the same methods as described above for TARGET cancer samples. Genes were retained if at least one sample had expression greater than 0 FPKM. To identify differentially expressed genes between siNTC and siTBX2-AS1 treated cells, we used the DESeq2 method with default parameters. Differentially expressed genes were annotated based on absolute value log fold change > 1.5 and Benjamini-Hochberg adjusted p-value < 0.1. Gene set enrichment analysis (GSEA) was performed across samples using the MsigDB Hallmarks gene sets and significantly enriched gene sets with FDR q-val < 0.1 were retained. Up-stream co-regulators of differentially expressed genes were identified using default parameters from the iRegulon program part of the Cytoscape suite.

### Data Availability

All TARGET RNA and DNA-sequencing data analyzed in this study are available through the database of Genotypes and Phenotypes (dbGaP; https://www.ncbi.nlm.nih.gov/gap/) under study-id phs000218 and accession number phs000467. GMKF RNA-sequencing data are available through dbGAP study accession phs001436.v1.p1. Neuroblastoma cell line RNA-sequencing data analyzed in this study are available through GEO at accessions GSE89413. NBL histone ChIP-seq and transcription factor ChIP-seq data used in this study are both available through GEO at accessions: GSE138315 and GSE94822, respectively. T-ALL transcription factor ChIP-seq data and SMC1 ChIA-PET data are available through GEO at accessions GSE29181, GSE59657, and GSE68977.

## Supporting information

Modi_Supplementary_Figures

Modi_Supplementary_Tables_1-3

Modi_Supplementary_Tables_4-6

Modi_Supplementary_Tables_7-9

Modi_Supplementary_Tables_10-12

Modi_Supplementary_Tables_13-15

Modi_Supplementary_Tables_16-18

## Acknowledgements

This work was supported in part by NIH grants R01-CA124709 (S.J.D.), R03 CA230366 (S.J.D.), X01 HL136997 (J.M.M.), and T32-HG46-18 (A.M.). This project was also funded in part by a supplement to the Children’s Oncology Group Chair’s grant CA098543 and with federal funds from the National Cancer Institute, National Institutes of Health, under Contract No. HHSN261200800001E to S.J.D and Complete Genomics. Promoter Capture C studies were funded by the Center for Spatial and Functional Genomics (A.D.W and S.FA.G) at CHOP. S.FA.G. was supported by the Daniel B. Burke Endowed and Chair for Diabetes Research and R01 HG010067.

## Author Contributions

A.M. and S.J.D. conceived and designed the study. M.A.S., J.M.G.A, D.S.G., E.J.P, S.M., S.P.H., S.J.D. and J.M.M. generated the TARGET data. K.L.C., M.E.J., S.J.D., A.D.W. and S.F.A.G. generated promoter-focused capture C data. E.M. C.S, and A.M. analyzed promoter-focused capture C data. A.M., G.L., S.R. analyzed TARGET data. A.M., K.L.C., T.C.L. and D.C. performed *TBX2-AS1* experiments. S.J.D. supervised the study. A.M. and S.J.D drafted the manuscript. All authors edited and approved the manuscript.

