## Supplementary material for "Integrative genomics identifies lncRNA regulatory networks across 1,044 pediatric leukemias and extra-cranial solid tumors": Modi_Supplementary_Figures

Apexa Modi^1,2^, Gonzalo Lopez^1,3^, Karina L. Conkrite^1^, Tsz Ching Leung^1^, Sathvik Ramanan^1^, Daphne Cheung^1^, Chun Su^4^, Elisabetta Manduchi^4^, Matthew E. Johnson^4^, Samantha Gadd^5^, Jinghui Zhang^6^, [Malcolm A. Smith](https://www.nature.com/articles/nature25795#auth-24)^7^, [Jaime M. Guidry Auvil](https://www.nature.com/articles/nature25795#auth-25)^8^, Daniela S. Gerhard^7^, Soheil Meshinchi^9^, Elizabeth J. Perlman^5^, Stephen P. Hunger^1,10^, John M. Maris^1,10,11^, Andrew D. Wells^4,12^, Struan F.A. Grant^4,13,14,15^, Sharon J. Diskin^1,10,11*^

^1^Center for Childhood Cancer Research, Children's Hospital of Philadelphia, Philadelphia, Pennsylvania 19104, USA.

^2^ Genomics and Computational Biology Graduate Group, Biomedical Graduate Studies, Perelman School of Medicine, University of Pennsylvania, Philadelphia, Pennsylvania 19104, USA.

^3^ Department of Genetics and Genomic Sciences, Icahn School of Medicine at Mount Sinai, New York 10029, USA.

^4^ Center for Spatial and Functional Genomics, Children's Hospital of Philadelphia, Philadelphia, Pennsylvania, USA.

^5^ Department of Pathology and Laboratory Medicine, Ann & Robert H. Lurie Children’s Hospital of Chicago, Robert H. Lurie Cancer Center, Northwestern University, Chicago, Illinois 60208, USA.

^6^ Department of Computational Biology, St Jude Children’s Research Hospital, Memphis, Tennessee 38105, USA

^7^ Cancer Therapy Evaluation Program, National Cancer Institute, Bethesda, Maryland 20892, USA.

^8^ Office of Cancer Genomics, National Cancer Institute, Bethesda, Maryland 20892, USA.

^9^ Clinical Research Division, Fred Hutchinson Cancer Research Center, Seattle, Washington 98109, USA.

^10^ Department of Pediatrics, Perelman School of Medicine at the University of Pennsylvania, Philadelphia, Pennsylvania 19104, USA.

^11^Abramson Family Cancer Research Institute, Perelman School of Medicine at the University of Pennsylvania, Philadelphia, Pennsylvania 19104, USA.

^12^Department of Pathology and Laboratory Medicine, Perelman School of Medicine at the University of Pennsylvania, Philadelphia, Pennsylvania 19104, USA.

^13^Department of Genetics, Perelman School of Medicine at the University of Pennsylvania, Philadelphia, Pennsylvania 19104, USA.

^14^ Division of Endocrinology & Diabetes, Children's Hospital of Philadelphia, Philadelphia, Pennsylvania, USA.

^15^ Divisions of Human Genetics, Children's Hospital of Philadelphia, Philadelphia, Pennsylvania, USA.

### **I. SUPPLEMENTARY TABLES**

**Supplementary Table 1: TARGET clinical sample and RNA-sequencing characteristics** **(xls file)** Overview of RNA-seq samples selected for final cohort and information about the type of RNA-sequncing available per cancer.

**Supplementary Table 2: Genomic loci for lncRNA and protein coding genes in this study (xls file)** Genomic position, gene type, HUGO gene name, and chromosomal band for all lncRNA and protein coding genes considered in this study.

**Supplementary Table 3: Number and types of genes expressed per cancer (xls file)** The number of protein coding genes and known and novel lncRNAs expressed per cancer after filtering out lowly expressed genes.

**Supplementary Table 4: Top 10 expressed lncRNAs across TARGET cancers and GTEx tissues (xls file)** The top 10 expressed lncRNAs for TARGET cancers and GTEX tissues ranked based on highest proportion of expression over the total sum of all lncRNA expression (FPKM).

**Supplementary Table 5: Tissue specificity index (tau score) annotation per gene (xls file)** The tissue specifcity index, calculated as the tau score, per gene. The cancer with the highest expression of that gene is also listed.

**Supplementary Table 6: Validation of tissue specific lncRNAs based on tau score analysis in alternate NBL datasets (xls file)** The number of tissue specific lncRNAs per cancer with NBL pure data set, which includes samples that were 80-90% free of immune/stromal cell infiltration and number of tissue specific lncRNAs per cancer with GMKF NBL dataset.

**Supplementary Table 7: Number of TARGET samples with WGS and SCNA events per cancer identified by GISTIC (xls file)** The number of TARGET cancers with available WGS and the number of samples that had matched RNA-seq and the SCNA events identified by GISTIC per cancer including number of samples per event.

**Supplementary Table 8: Differential expression of genes in samples with and without SCNA** **(xls file)** List of all lncRNAs in SCNA regions and their log2 fold change and p-value (Wilcoxon rank sum test) between samples with and without SCNA.

**Supplementary Table 9: Number of samples with SVs in lncRNAs (xls file)**

List of all lncRNAs and number of samples of the specific cancer with a structural variant breakpoint in or near (+/- 2.5kb) the lncRNA and annotation of whether the lncRNA is located in an SCNA region of that cancer.

**Supplementary Table 10: Statistics for input and output variables of lncMod analysis (xls file)** Input parameters for lncMod analysis including the number of dysregulated lncRNAs, expressed transcription factors, and significant TF-target gene associations per cancer. The proportion and number of significantly dysregulated lncMod triplets compared to the number of all possible triplets per cancer.

**Supplementary Table 11: Significantly dysregulated lncMod triplets (xls file)** List of all significantly dysregulated lncMod triplets, which include a lncRNA modulator, transcription factor, and target gene.

**Supplementary Table 12: lncRNA TF associations ranked by # target genes (xls file)** The top 10 transcription factors, associated with a lncRNA modulator, ranked based on number of target genes associated with the transcription factor and lncRNA modulator per cancer.

**Supplementary Table 13: lncRNAs associated with CRC of T-ALL/NBL (xls file)** lncRNAs identified to be regulated by the CRC of either T-ALL, NBL, or both independently.

**Supplementary Table 14: Correlation between CRC regulated lncRNAs and PCGs. Gene set enrichment analysis of correlated CRC PCGs (xls file)** CRC regulated lncRNAs and protein coding genes that have significant expression correlation with each other in either NBL or T-ALL and gene set enrichment results for CRC regulated protein coding genes with significant correlation to a CRC regulated lncRNA. Gene set enrichment analysis performed with Fisher exact test and MsigDB hallmark gene sets.

**Supplementary Table 15: Differentially expressed lncRNAs between major subtypes in NBL and T-ALL (xls file)** Differential expression analysis results comparing lncRNA expression between MYCN amplified and non-amplified NBL samples. Differential expression analysis results comparing lncRNA expression between the TAL1 subgroup and other T-ALL sample subgroups. lncRNAs that are differentially expressed in both NBL and T-ALL and annotation of whether the lncRNA is regulated by CRC transcription factors.

**Supplementary Table 16: Data integration using multi-dimensional analysis to prioritize functional lncRNAs in each cancer (xls file)** lncRNAs prioritized as likely functional based on association with a particular cancer through the results of various analyses used in this study.

### **Supplementary Table 17: Prioritized lncRNAs and predicted lncRNA target genes and pathways in NBL (xls file)** lncRNAs prioritized as likely functional based on all analyses used in this study.

### **Supplementary Table 18: GSEA Analysis: MsigDB Hallmarks enriched across genes impacted by siTBX2-AS treatment in NLF (xls file)** Results from GSEA of genes significantly up- or down-regulated due to siTBX2-AS1 treatment in the neuroblastoma cell line: NLF.

### **II. SUPPLEMENTARY FIGURES**

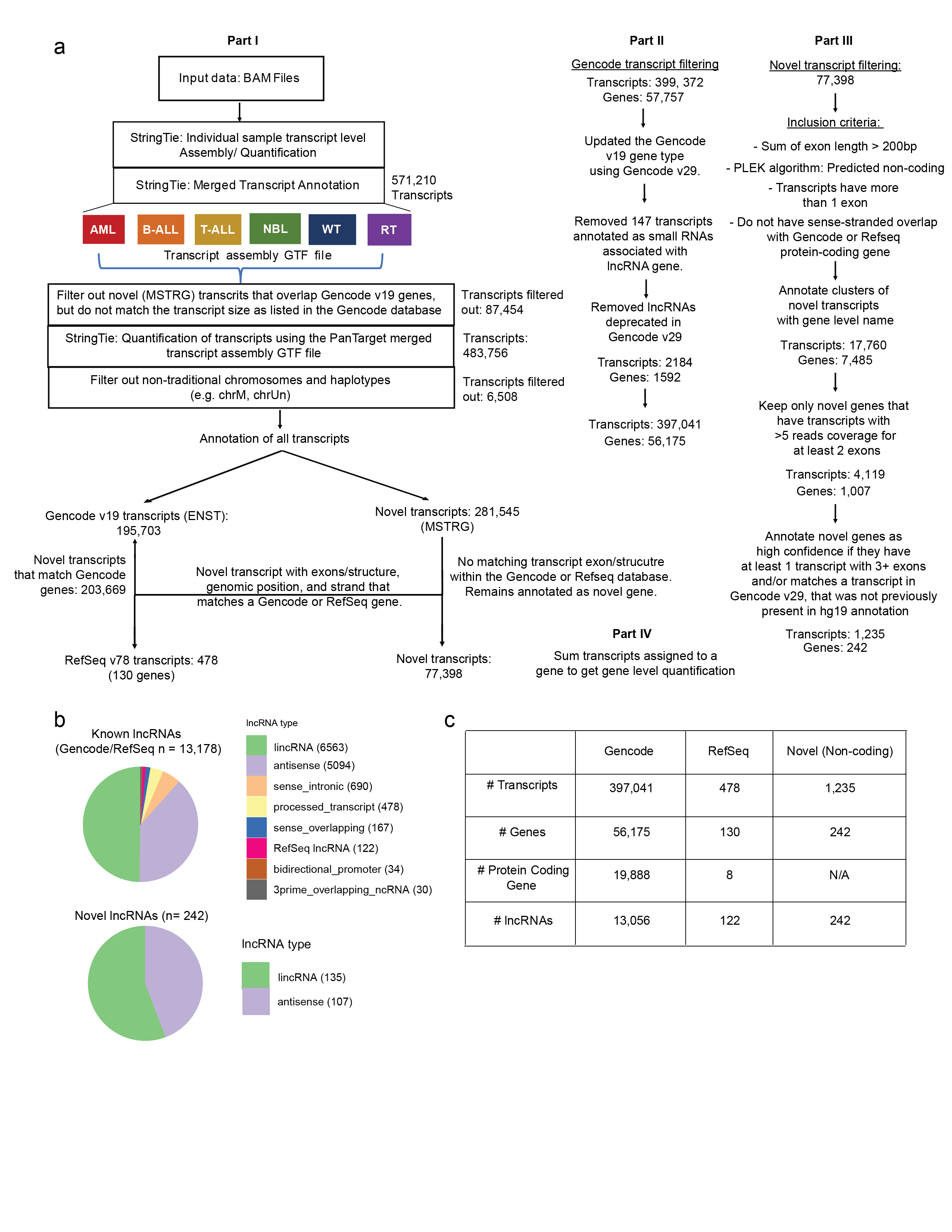

**Supplementary Figure 1: Workflow for RNA-seq gene mapping and quantification.** (a) Workflow diagram showing how samples were processed using the StringTie program, which performs genes mapping and quantification. Custom scripts were then used for the following: Part I: Identified gene transcripts were assigned as a Gencode or Refseq gene, or as a novel gene. Part II: Gencode transcripts were further filtered including based on gene type. Part III: Novel genes were further filtered based on non-coding potential, length, number of transcripts and exons, and read coverage per exon. Part IV: Gene-level expression was considered as the sum of associated transcript expression. (b) High confidence selected novel lncRNA genes are primarily intergenic or antisense. Sense-overlapping novel lncRNAs were considered low confidence and not-considered in this analysis. The majority of known lncRNAs in the Gencode database are either intergenic or antisense. (c) The number of transcripts and genes per gene type post filtering used in this study.

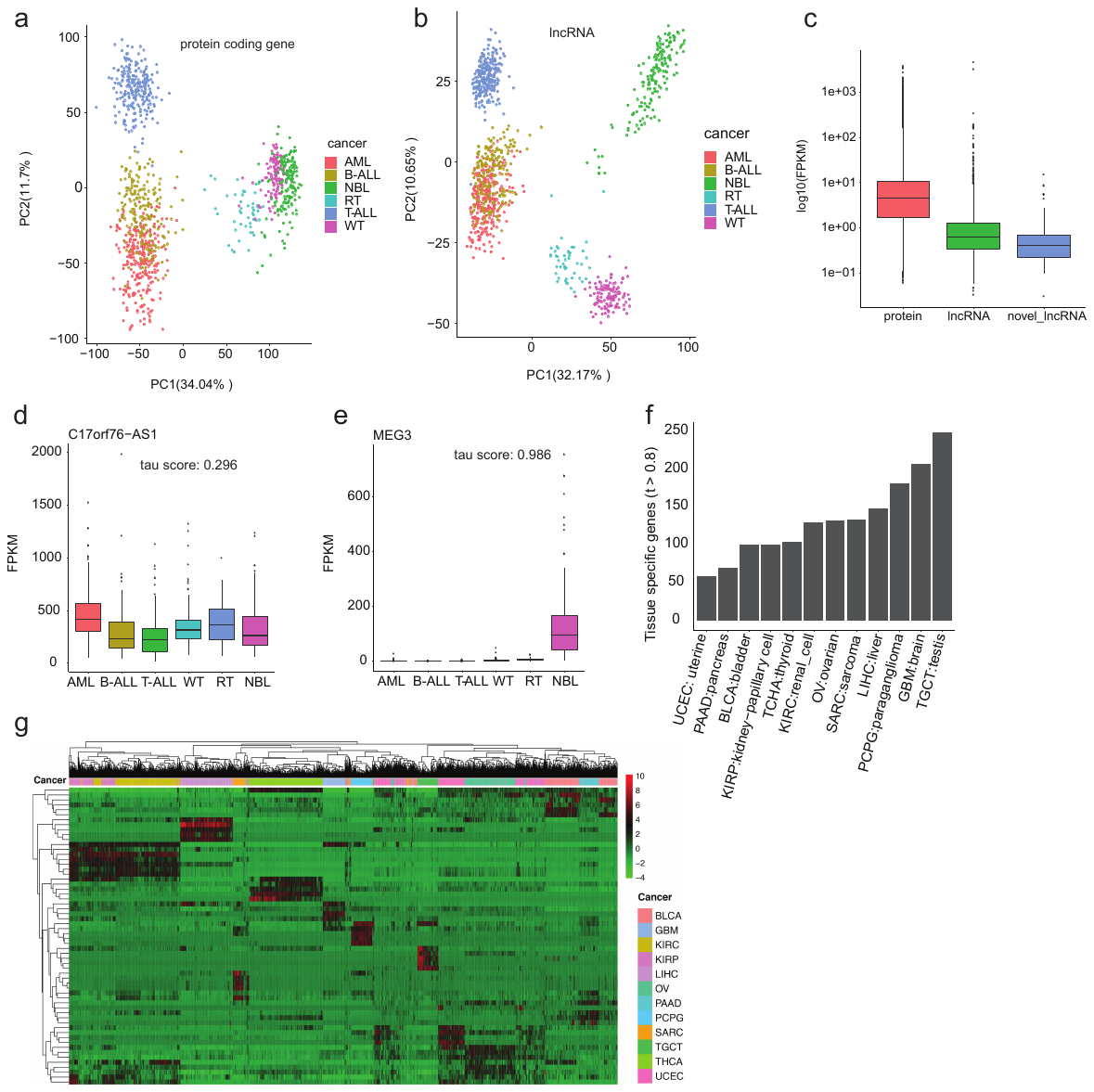

**Supplementary Figure 2: lncRNA expression varies across pediatric cancers.** (a) Average expression of protein coding genes, known, and novel lncRNAs in order of highest average expression. (b) PCA of protein coding gene expression across pediatric cancers. (c) PCA showing unsupervised clustering of lncRNA gene expression alone. Each cancer is more closely clustered together individually than in the PCA including protein coding genes, suggesting that each cancer has distinct lncRNA expression, except for AML and B-ALL, which appear to have very similar lncRNA expression. (d) Expression of a ubiquitously expressed lncRNA: *C17orf76*-*AS1* and its tau score: 0.296 is low. (e) Expression of the *MEG3* lncRNA is primarily in NBL and thus has a higher tau score of 0.986, (tau score >0.8 indicates tissue specificity). (f) The number of tissue specific lncRNAs across 12 adult cancers from TCGA. (g) Unsupervised clustering of the expression of the top 5 most TS lncRNAs (ranked by expression and tau score) in 12 adult cancers (total lncRNAs n=60).

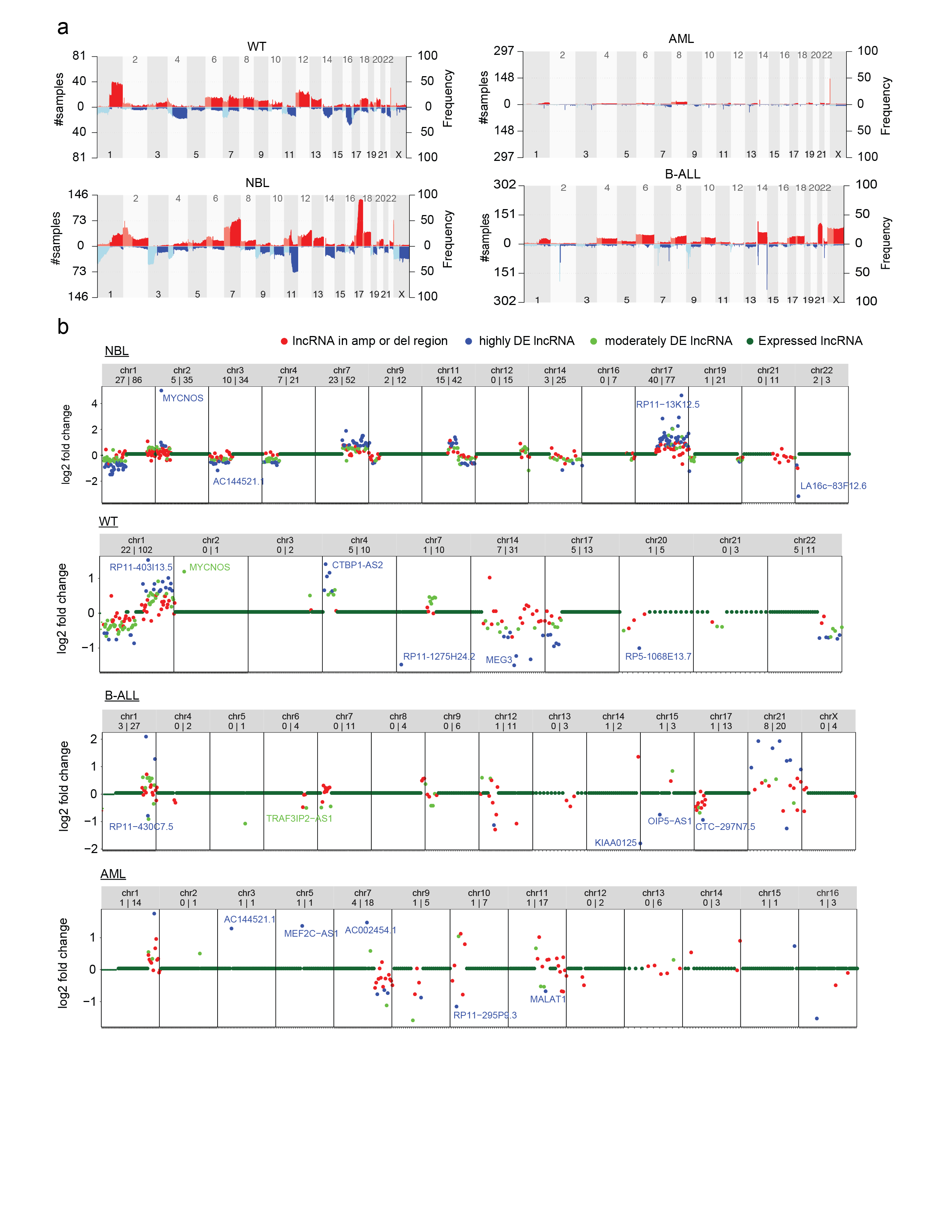

**Supplementary Figure 3: Regions of somatic copy number aberration across cancers and genes dysregulated due to copy number.** (a) Plots of the frequency of copy number gain and loss across the genome for four pediatric cancer cohorts in this study. (b) lncRNA loci on chromosomes with copy number alterations across the pediatric cancers: NBL, WT, B-ALL, and AML. lncRNAs were evaluated to have differential expression due to copy number using the Wilcoxon rank sum test: highly differential: p-value < 0.05 and log |fold change| > 1.5 and moderately differential: p-value < 0.1 and log |fold change| > 1.0. Points are colored based on loci in an amplified or deleted region of the chromosome and if the lncRNA is highly or moderately differentially expressed.

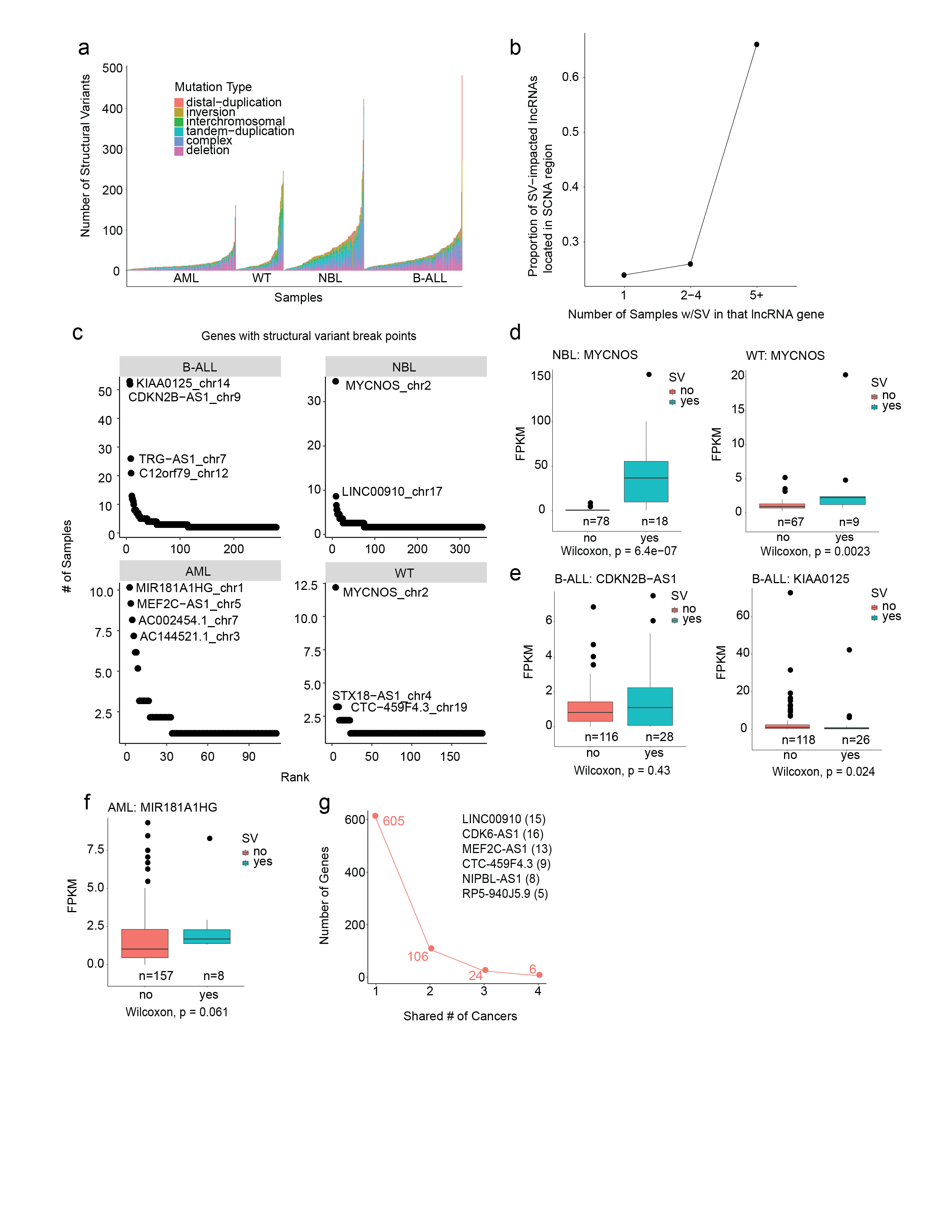

**Supplementary Figure 4: Structural variants impact lncRNAs in pediatric cancers.** (a) The number and types of structural variants as annotated by Complete Genomics (CGI) per sample per cancer. (b) The number of samples with a structural variant in a lncRNA or protein coding gene (gene-sample pairs) vs number of lncRNA genes also found in copy number regions. (c) Ranking of genes with structural variants by the number of samples per cancer. (d) Expression of the top genes per cancer in samples with and without structural variant (NBL/WT: MYCNOS, (e) B-ALL: CDKN2B-AS1,KIA00125, and (f) AML: MIR18A1HG. (g) The number of genes with a structural variant found in 1-4 cancers.

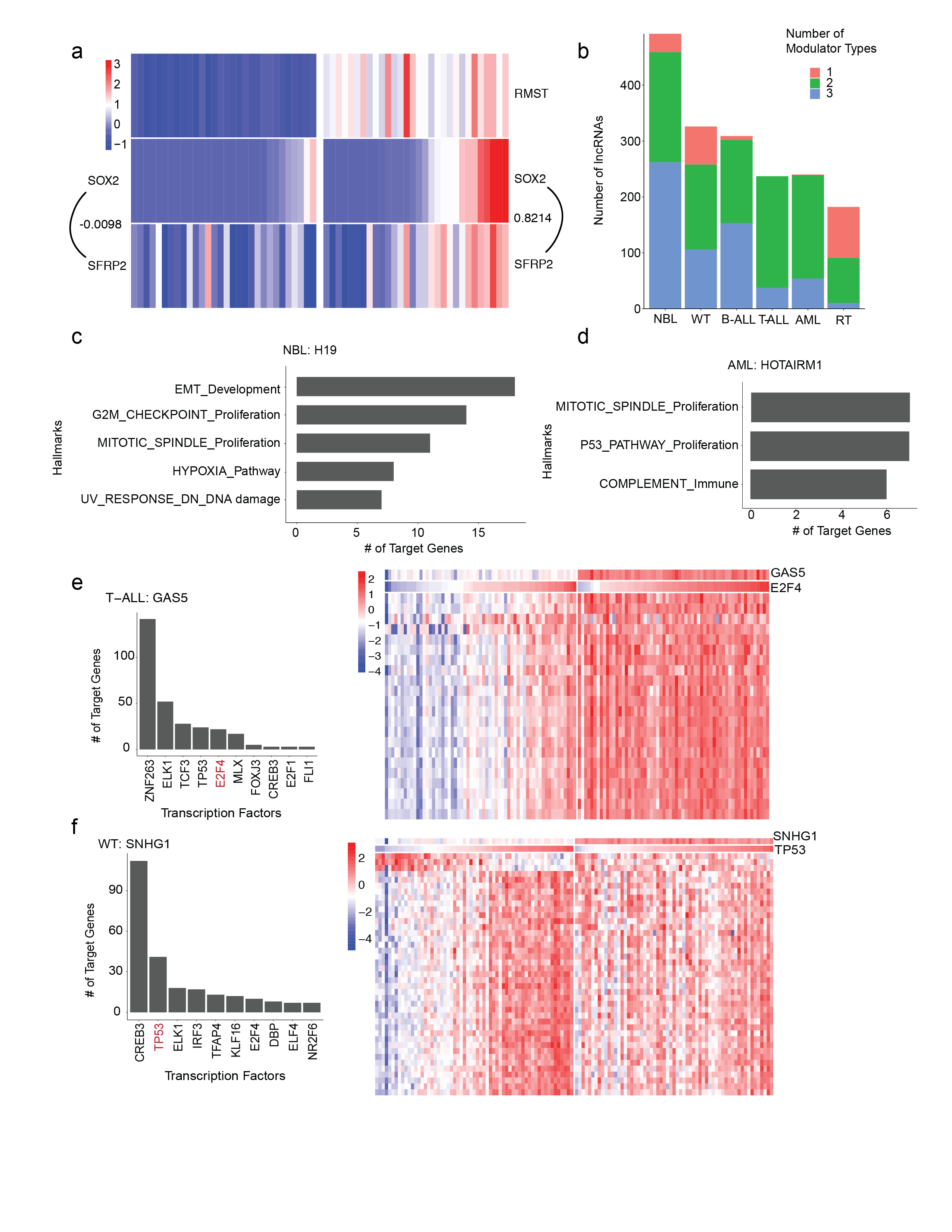

**Supplementary Figure 5: Characteristics of lncMod analysis and results.** (a) Example of a dysregulated lncMod triplet in Wilm’s Tumors (WT). Samples with high *RMST* expression also have higher expression correlation between *SOX2* and its target gene *SFRP2*. SOX2 regulation of *SFRP2* appears to be disrupted in samples with low *RMST* expression, as suggested by the low expression correlation of TF and target gene in these samples. (b) The number of modulator types (attenuate, enhance, or invert) associated with a lncRNA modulator based on its impact of a specific TF-target gene. (c) Number of target genes of the *H19* lncRNA modulator in NBL that are enriched in MsigDB hallmarks gene sets. (d) Target genes of the *HOTAIRM1* lncRNA modulator in AML that are enriched in MsigDB hallmarks gene sets. (e) The top transcription factors, based on number of associated dysregulated target genes, impacted by the *GAS5* lncRNA modulator in T-ALL and the expression of *GAS5*, the transcription factor *E2F4* and its target genes in T-ALL samples with *GAS5* expression dysregulation. (f) The top transcription factors, based on number of associated dysregulated target genes, impacted by the *SNHG1* lncRNA modulator in WT and the expression of *SNHG1*, the transcription factor *TP53* and its target genes in WT samples with *SNHG1* expression dysregulation.

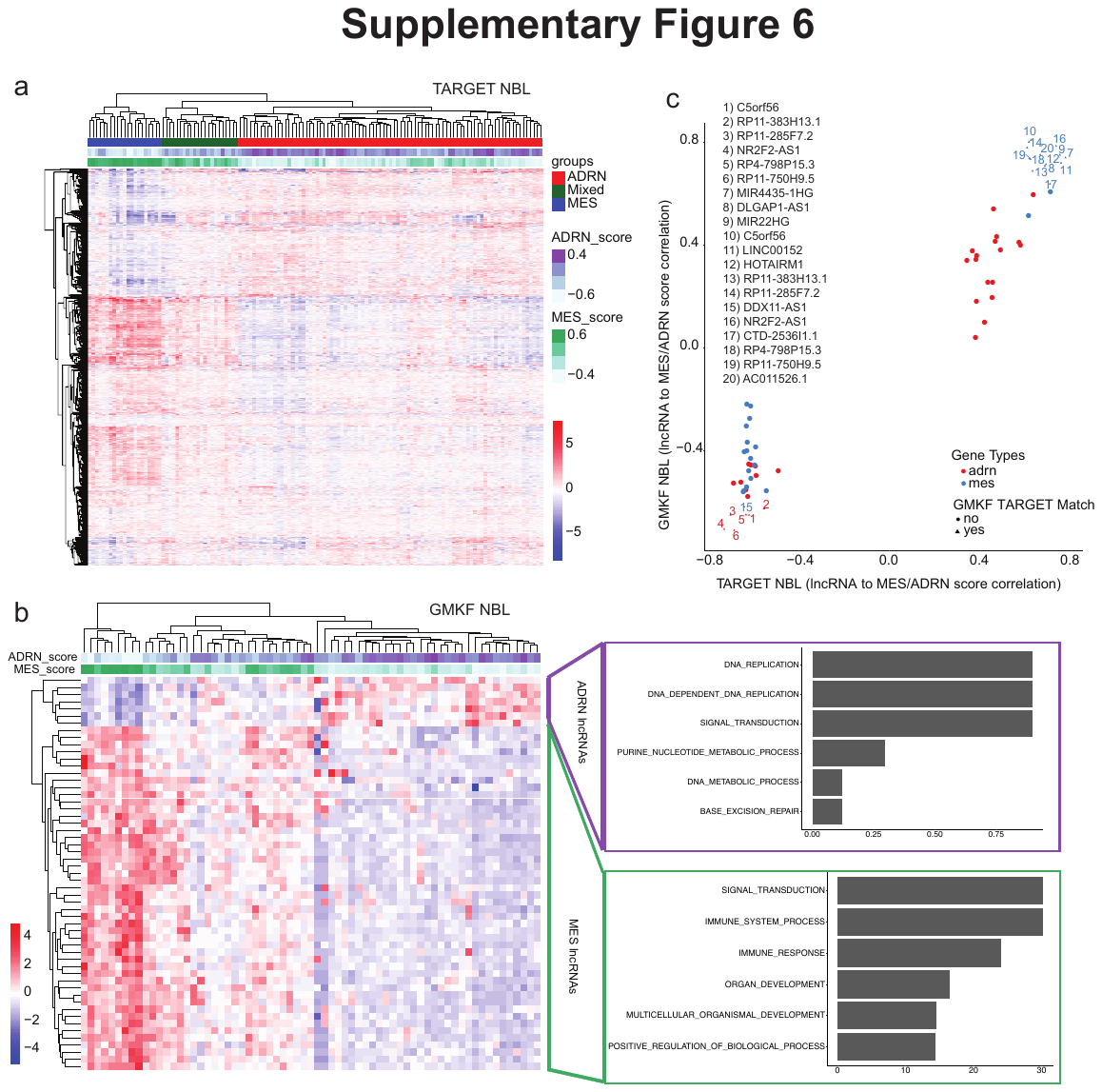

**Supplementary Figure 6: Defining mesenchymal vs adrenergic lncRNAs in two NBL cohorts.** (a) Heatmap of the expression of genes previously shown to be associated with either the MES or ADRN cell lineage in the TARGET NBL cohort. Samples were assigned into three groups using hierarchical clustering based on whether they had more expression of MES or ADRN genes. (b) Expression of lncRNAs with significant correlation (|r| >0.6) to the MES or ADRN score in the GMKF NBL cohort. lncRNAs were then correlated with protein coding genes on the same chromosome and subsequent gene set enrichment analysis was performed for MES and ADRN protein coding genes separately. Gene set enrichment results for each group are shown to the right of the heatmap. (c) Correlation of lncRNAs to the MES and ADRN score in the TARGET (x-axis) and GMKF (y-axis) NBL cohort respectively. Numbered points represent lncRNAs that had a significant correlation to the MES or ADRN score in both cohorts.

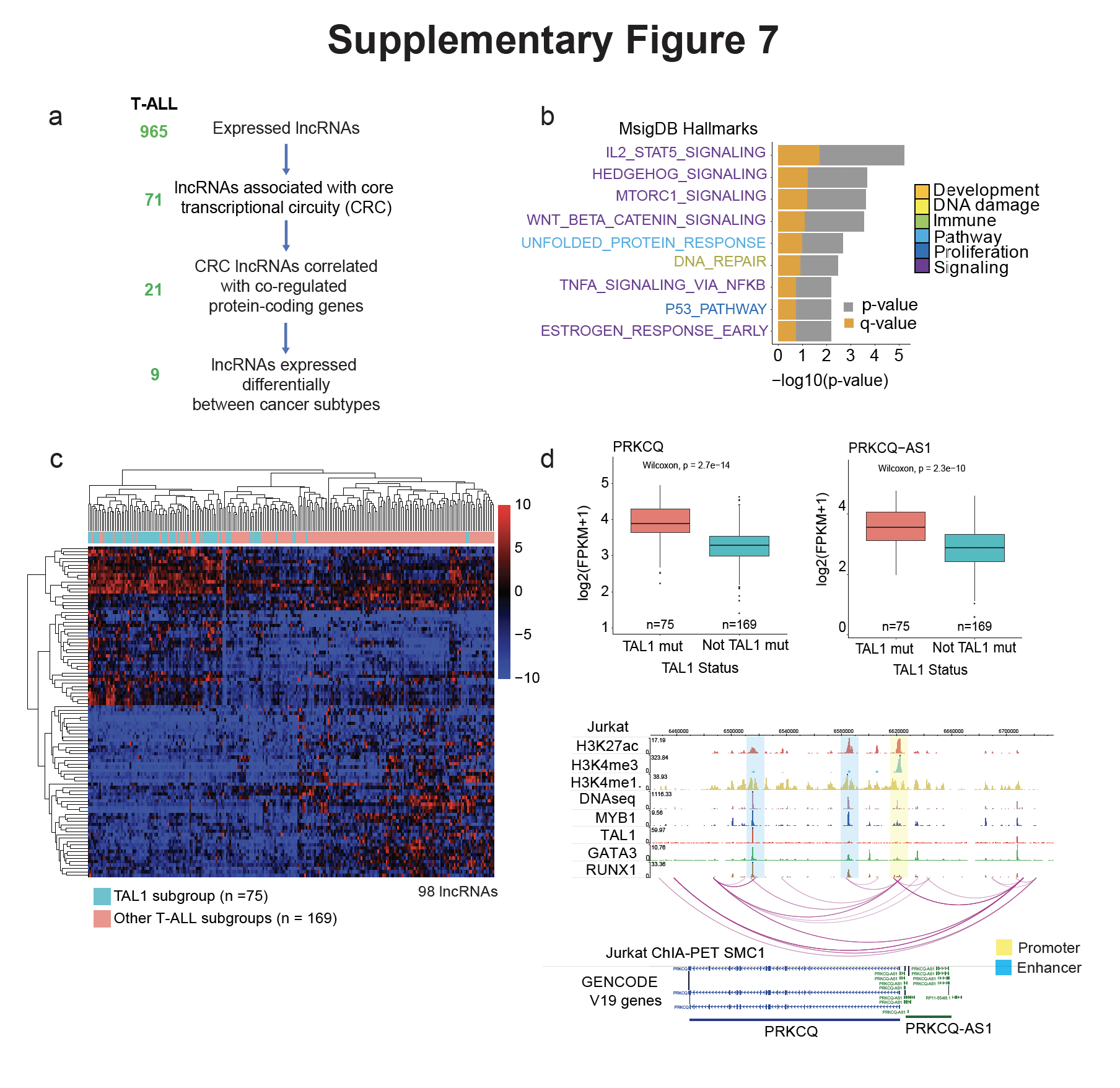

**Supplementary Figure 7: Identification of cancer associated lncRNAs regulated by core transcription factors in T-ALL.** (a) Filtering of lncRNAs expressed in either NBL or T-ALL based on CRC TF regulation, differential expression based on cancer subtypes, and co-regulation with a CRC associated protein-coding gene. In T-ALL, differentially expressed lncRNAs are between the T-ALL cancer subtypes: TAL1 subgroup vs non-TAL1 subgroup. Co-regulation of CRC regulated lncRNAs and protein coding genes was determined by correlation analysis. (b) Gene set enrichment analysis results for protein coding genes significantly correlated with CRC regulated lncRNAs (Pearson’s r > 0.4 and FDR < 0.1). (c) Heatmap of the lncRNAs that are differentially expressed between the TAL1 subgroup vs non-TAL1 subgroup (d) Expression of *PRKCQ* and *PRKCQ*-*AS1* stratified by T-ALL subgroups. ChIP-seq tracks for histone marks and CRC transcription factors and ChIA-PET chromatin interactions in the T-ALL cell line: Jurkat at the *PRKCQ*/ *PRKCQ*-*AS1* locus.

**Supplementary Figure 8: Validation of *TBX2*-*AS1* expression in NBL cell lines and impact on NBL cell growth.** (a) Expression of *TBX2-AS1* and *TBX2* across pediatric cancers. (b) Correlation between *TBX2-AS1* and other genes known to be regulated by TBX2. (c) Expression of *TBX2-AS1* and *TBX2* in 38 NBL cell lines identified from publicly available RNA-sequencing. (d) RT-qPCR validation of *TBX2*-*AS1* and *TBX2* expression in 8 NBL cell lines. (e) Volcano plot showing genes with significant dysregulation (log fold change > 1.5 and -log10 p-value <0.1) observed in transcriptomic profiling of NLF cells treated with siTBX2-AS1. (f) Representative image of cell growth (as measured by RT-Ces assay) of the NBL cell lines: NLF and SKNSH with all treatments. Cell index is normalized to time point when siRNA reagent is added at 24 hours post cell plating. (g) Images of the NBL cell line: NLF, which has the highest expression of *TBX2*-*AS1*, 72 and 96 hours post-transfection with siRNAs targeting *TBX2*-*AS1* and non-targeting control (siNTC).
